# Conserved, divergent and heterochronic gene expression during *Brachypodium* and *Arabidopsis* embryo development

**DOI:** 10.1101/2021.03.05.434107

**Authors:** Zhaodong Hao, Zhongjuan Zhang, Daoquan Xiang, Prakash Venglat, Jinhui Chen, Peng Gao, Raju Datla, Dolf Weijers

## Abstract

Embryogenesis, transforming the zygote into the mature embryo, represents a fundamental process for all flowering plants. Current knowledge of cell specification and differentiation during plant embryogenesis is largely based on studies of the dicot model plant *Arabidopsis thaliana*. However, the major crops are monocots and the transcriptional programs associated with the differentiation processes during embryogenesis in this clade were largely unknown. Here, we combined analysis of cell division patterns with development of a temporal transcriptomic resource during embryogenesis of the monocot model plant *Brachypodium distachyon*. We found that early divisions of the *Brachypodium* embryo were highly regular, while later stages were marked by less stereotypic patterns. Comparative transcriptomic analysis between *Brachypodium* and *Arabidopsis* revealed that the early and late embryogenesis shared a common transcriptional program, whereas mid-embryogenesis was divergent between species. Analysis of orthology groups revealed widespread heterochronic expression of potential developmental regulators between the species. Interestingly, *Brachypodium* genes tend to be expressed at earlier stages than *Arabidopsis* counterparts, which suggests that embryo patterning may occur early during *Brachypodium* embryogenesis. Detailed investigation of auxin-related genes shows that the capacity to synthesize, transport, and respond to auxin is established early in the embryo. However, while early PIN1 polarity could be confirmed, it is unclear if an active response is mounted. This study presents a resource for studying *Brachypodium* and grass embryogenesis, and shows that divergent angiosperms share a conserved genetic program that is marked by heterochronic gene expression.

**Key message:** Developmental and transcriptomic analysis of *Brachypodium* embryogenesis, and comparison with *Arabidopsis*, identifies conserved and divergent phases of embryogenesis, and reveals widespread heterochrony of developmental gene expression.

## Introduction

Angiosperms represent a diverse group of plants that share a number of characteristics: a dominant diploid sporophytic state, true embryos with precursors for the major tissues, including meristems, an elaborate vascular transport system, seeds and flowers. Both major groups of angiosperms: dicots and monocots, encompass crops as well as genetic model organisms. In both groups, the embryo represents a relatively simple form in which – from a fertilized egg cell – a miniature plant emerges that has primordial organs and tissues, including meristems that sustain post-embryonic growth. Few models have been used to extensively study progression and genetics of embryo development, and these include the dicots tobacco, *Arabidopsis thaliana*, and soybean, as well as the monocots rice, maize, and wheat (Armenta-Medina et al. 2021; Palovaara et al. 2016). From these analyses, as well as from earlier comparative embryology (Johri 1984), it is evident that the morphology and developmental progression is very different between dicots and monocots. In fact, it is difficult to even identify homologous stages based on morphology. Thus, whereas there is a prominent body of literature on genetic regulation of *Arabidopsis* embryogenesis (Palovaara et al. 2016; ten Hove et al. 2015), it is far from trivial to transpose this towards monocot plants (Zhao et al. 2017). Following the identification of developmental regulators in *Arabidopsis*, analysis of expression patterns of maize or rice homologs have shown that there is both conservation and divergence of expression patterns. For example, within the *WOX* family, some members show different patterns between *Arabidopsis* and maize (Haecker et al. 2004; Nardmann et al. 2007), but the pattern of *WOX5* appears conserved between *Arabidopsis*, maize, and rice (Kamiya et al. 2003; Nardmann et al. 2007; Sarkar et al. 2007). Likewise, the *Arabidopsis STM* and maize *KN* genes have similar expression (Kerstetter et al. 1997; Long and Barton 1998; Smith et al. 1995). Thus, a major open question is how (dis)similar embryo developmental patterns and their regulation are between monocots and dicots.

Several studies have focused on monocot embryogenesis from either a morphological (Black et al. 2006; Guillon et al. 2012; Itoh et al. 2005; Smart and O’Brien 1983) or transcriptional (Chen et al. 2017; Itoh et al. 2016; Yi et al. 2019) perspective. From these however, it is not yet clear how the developmental transitions and emergence of pattern elements are connected to genome-wide gene expression patterns. At the same time, it is not yet clear how the transcriptional landscape of monocot embryogenesis relates to that found in dicots.

Here, we focus on the development of the *Brachypodium distachyon* embryo. *Brachypodium* is a monocot grass model plant (Scholthof et al. 2018) that is closely related to wheat, yet is diploid, has a small size and short life cycle that allows cultivation in lab conditions, and has not been domesticated. Thus, it represents a “wild” grass model. The *Brachypodium* genome has been sequenced (The International Brachypodium Initiative. 2010), and the species is being used as model for bioenergy (Cass et al. 2016), root development (Agapit et al. 2020) and flowering (Qin et al. 2017), among others. Given that closely related crop relatives, such as wheat, are seed crops, there is an interest in understanding the control of embryo and grain development. An initial description of grain development (Guillon et al. 2012) showed that general developmental patterns of embryo development are comparable between *Brachypodium* and other grasses.

Here, we combined detailed analysis of cell division patterns with stage-specific transcriptome analysis to provide insights into *Brachypodium* embryogenesis. By comparative transcriptomics, we find that there early and late embryo phases share genetic programs between *Brachypodium* and *Arabidopsis*, whereas mid-embryogenesis is divergent. Analysis of orthology groups reveals widespread heterochrony of embryo development, where *Brachypodium* appears to express many genes at earlier stages than the *Arabidopsis* counterpart. Detailed investigation of auxin transport and response shows conserved expression between species, but it is unclear if the hormone controls embryogenesis in *Brachypodium*. Thus, embryogenesis in *Brachypodium* is marked by a conserved angiosperm transcriptional program, as well as lineage-specific programs and heterochronic expression of many potential regulators.

## Results

### *Brachypodium distachyon* embryo development

Embryo development of several grass species have been described. Generally, embryo stages are comparable between *Brachypodium* and wheat (Guillon et al. 2012; Xiang et al. 2019). Here, we extended earlier descriptions of embryogenesis with an emphasis on early, morphogenetic stages. Through whole-mount microscopy and scanning electron microscopy, we confirmed the previously described stages, and here systematically name these as two-cell embryo or quadrant (TCQ; Fig. 1A), pre-embryo (PEM; Fig. 1B), transition (TRA; Figs. 1C, D), leaf early (LEE; Figs. 1E-G), leaf middle (LEM; Fig. 1H, I), leaf late (LEL; Fig. 1J, K), and mature (MAT; Fig. 1L, M). For the earliest stages, we additionally performed ClearSee-based staining (Ursache et al. 2018), followed by high-resolution confocal microscopy (Fig. 1N-U) and cell segmentation (Fig. 2) (Yoshida et al. 2014). In the following, we describe the morphogenetic hallmarks of embryo progression and its cellular basis.

**Figure 1.**
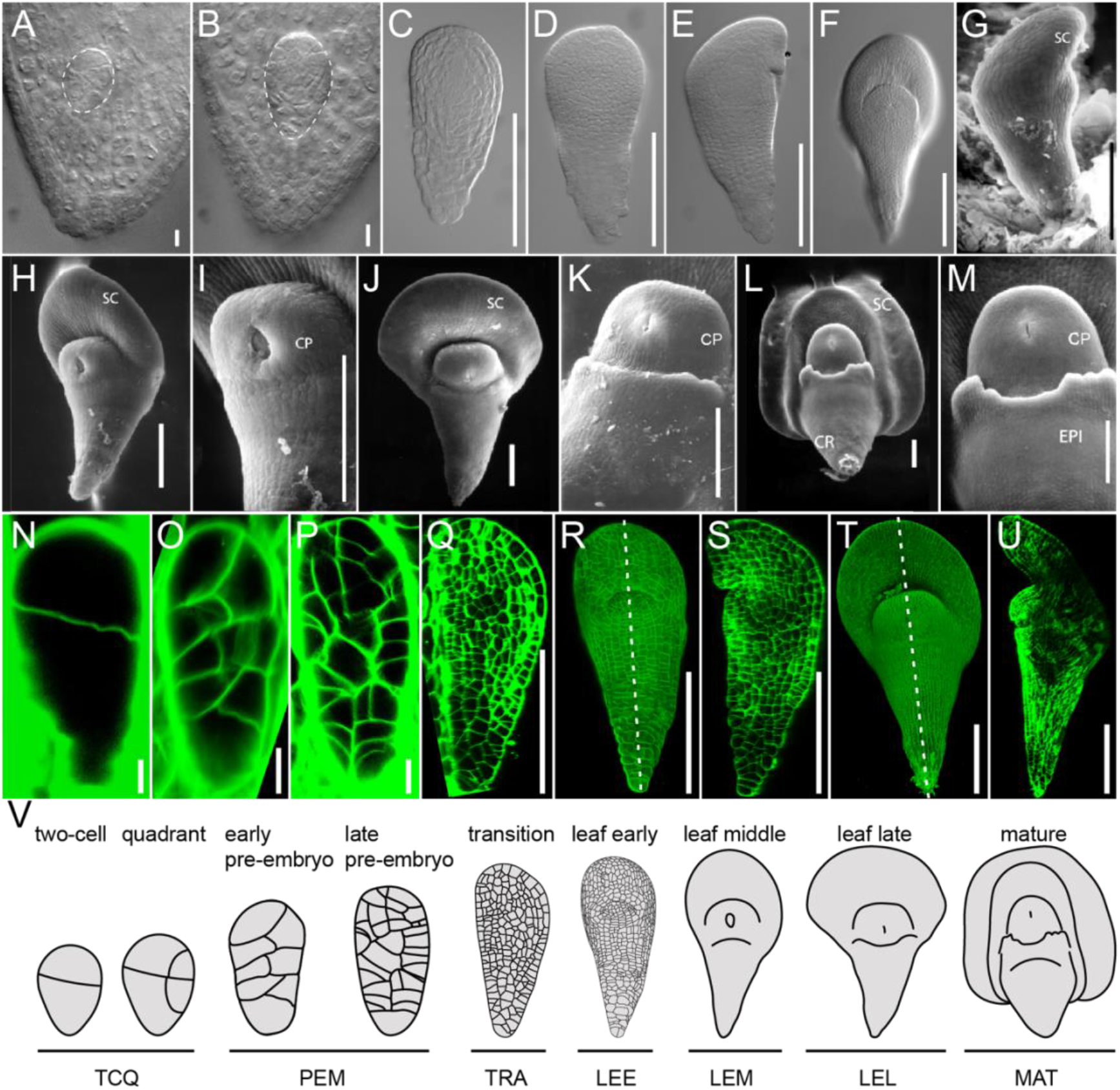
Development of *Brachypodium* embryos. Development in successive stages of Brachypodium embryos was visualized by light microscopy (a-f), scanning electron microscopy (g-m), and confocal imaging (n-u). Stages are two-cell or quadrant (a, n), pro-embryo (b, o, p), transition (c, d, q), leaf early (e-g, r, s), leaf middle (h, i, t, u), leaf late (j, k), and mature (l, m). (s, u) are optical section along the white dashed lines in (r, t). Embryos in (a, b, n-p) are inside seeds, while all others were removed from the seed. (v) Full series of developmental stages and nomenclature. Scale bars: 5 μm in (a, n), 10 μm in (b, o, p), 50 μm in (i, k, m) and 100 μm in (c-h, j, l, q-u). SC, scutellum; CP, coleoptile; CR, coleorhiza; EPI, epiblast.

**Figure 2.**
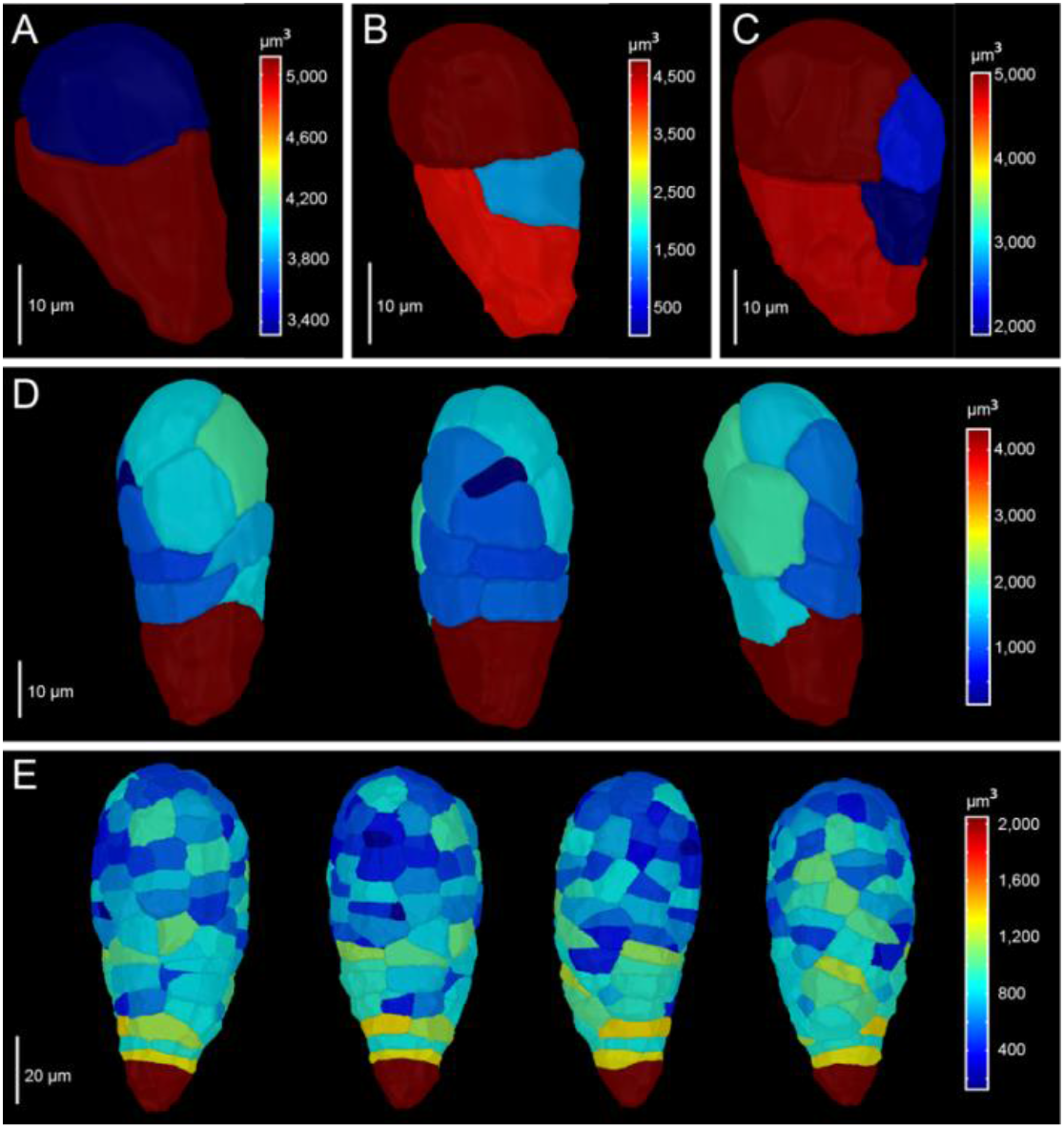
Division patterns in early Brachypodium embryos. *Brachypodium* embryos were imaged and segmented at two-cell (a), three-cell (b), four-cell (c), pre-embryo (d), and transition stages (e) by using confocal microscopy and MorphographX. In (d) and (e), the same embryos is shown from different angles. False color scale shows the volume of cells.

Analysis of the earliest stages of embryogenesis revealed that zygote division is asymmetric (Fig. 1N & Fig. 2A), generating a small apical and large basal cell. Following this initial, asymmetric division, each of the daughter cells again divides asymmetrically, generating two neighboring small cells (Fig. 2B, C). While the following divisions were less stereotypical, the pattern of divisions generated a cluster of small cells, likely from the initial two small daughter cells, surrounded by a group of larger cells (Fig. 2D, E). At early stages, the embryo thus already displays the laterally bent structure that characterizes later stages. From the lateral region of larger cells, the scutellum originates, while a dome that marks the shoot apical meristem (SAM) area arises underneath the scutellum (Fig. 1R, S, V). An axis of smaller cells extends basally from the SAM, but distinct tissues, such as the future root meristem, are not readily anatomically recognizable until the leaf late stage (Fig. 1U). The scutellum was developed into a shield-shape, and a bulging coleoptile was clearly observed in the leaf middle stage (Fig. 1T, U, V). At the same time, differentiation of epiblast cells occurred and further developed in the two subsequent stages, the leaf late and mature stages (Fig. 1J-M, V). Thus, while clear organs and structures are anatomically visible at the LEE, LEM, LEL, and MAT stages, no clear landmarks of patterning can be observed prior to this. Yet, the first divisions of the zygote are regular, which suggests the potential existence of an early pattern formation process.

### A reference transcriptome of the developing *Brachypodium* embryo

To generate molecular insight in the developmental progression of *Brachypodium* embryogenesis, we sampled isolated embryos from the 7 stages discussed in the previous section. Considering the substantial morphological changes occurring between the leaf early stage and the leaf middle stage, we collected an additional embryo stage, with an embryo length of 180 ± 25 μm (E180), between these two stages. We also collected embryo samples with a length of 400 ± 25 μm (E400), showing micro-morphological characteristics that were identical to that of embryos at the LEM stage. In addition to these 9 embryo stages, we also collected three non-embryo tissues, including early endosperm (EEN), late endosperm (LEN), and seed coat (SEC), corresponding to the embryo stage of TRA, LEL, and LEL, respectively. These were sampled with the aim to generate a reference transcriptome to correct for contamination with abundant endosperm and seed coat tissues during embryo isolation.

We next performed RNA sequencing (RNA-Seq) analysis on duplicates or quadruplicate of all 9 embryo stages and 3 endosperm and seed coat samples. Among the 34,260 annotated protein-coding genes in the *Brachypodium* genome (The International Brachypodium Initiative. 2010), 69.32% (23,749 genes) were expressed in at least one sample and 66.99% (22,951 genes) were differentially expressed (FDR < 0.05) between at least two different samples. Hierarchal clustering of the significant changes in gene expression across all samples revealed a progressive transcriptomic shift during *B. distachyon* embryo development and an obvious tissue-specific transcriptome profile between embryo and non-embryo tissues (Supplementary Figs. 1 & 2).

A major concern with manual dissection and sampling of early embryos from the much larger seed coat and dense endosperm is the contamination with non-embryo tissues. Studies in *Arabidopsis* showed that such contamination may confound embryo transcriptome profiling, and lead to contentious inferences (Schon and Nodine 2017). Thus, despite making great efforts to avoid contamination during embryo sampling (see Methods), we quantified the expression of some well-known tissue-specific genes across all samples to address the degree of non-embryo tissue contamination. Glutelin is a well-known seed storage protein, which has an endosperm-specific expression pattern in rice (Takaiwa et al. 1996). In the *B. distachyon* genome, glutelin is encoded by seven genes, all of which are specifically expressed in endosperm (Supplementary Fig. 3). Cellulose biosynthesis plays a very important role during seed coat development, particularly in secondary cell wall reinforcement and mucilage attachment (Griffiths and North 2017; Mendu et al. 2011). Seven cellulose synthases are expressed during *Brachypodium* embryogenesis, four of which are highly expressed in seed coat (Supplementary Fig. 4). In addition, studies in *Arabidopsis* (Kunieda et al. 2013), soybean (Gijzen et al. 1993) and prickly sida (Egley et al. 1983) showed that some peroxidases, heme-containing proteins, accumulated in, and contribute to seed coat development. Among the 154 peroxidases in the *Brachypodium* genome, 82 are expressed during *Brachypodium* embryogenesis and 17 are highly expressed in seed coat (Supplementary Fig. 5). None of these inferred endosperm- or seed coat-enriched transcripts was found to be expressed at appreciable levels in the isolated embryos (Supplementary Figs. 3-5), and indeed, principal component analysis clearly separates the endosperm and seed coat samples from all embryo samples (Supplementary Fig. 2). We thus conclude that there is minimal contamination in the embryo samples.

### Cross-species genomic conservation of embryo development

Morphological patterns of embryo development are very different between monocotyledonous and dicotyledonous plants, and it is an entirely open question whether the progression of developmental events and biological processes follows similar or different trajectories between these divergent groups. To address this question, we used our high-quality temporal transcriptome series for a comparison with datasets derived from the dicot *A. thaliana*, for which several embryo transcriptomes have been reported.

We combined two datasets (Hofmann et al. 2019; Nodine and Bartel 2012) to cover both early embryogenesis and late embryogenesis at high temporal resolution. A total of 19,893 genes (41.14% of all annotated genes) could be detected in embryo samples, and 17,314 genes are differentially expressed (FDR < 0.05) between any two of the developmental stages. A principal component analysis (PCA) showed that the first two PCs cumulatively explained 69.23% of the total variance and all samples were separated according to their developmental stage (Fig. 3B). We generated a comparable PCA plot for our *Brachypodium* dataset. The PC1 and 2 represented 73.26% of the total variance and stratified all samples in a successive, but distinct developmental trajectory (Fig. 3A). The PCA analyses indicate that although great differences exist in the external embryo development of *A. thaliana* and *B. distachyon*, both of their embryogenesis appear a gradual development not only at the morphological level but also at the global gene expression level.

**Figure 3.**
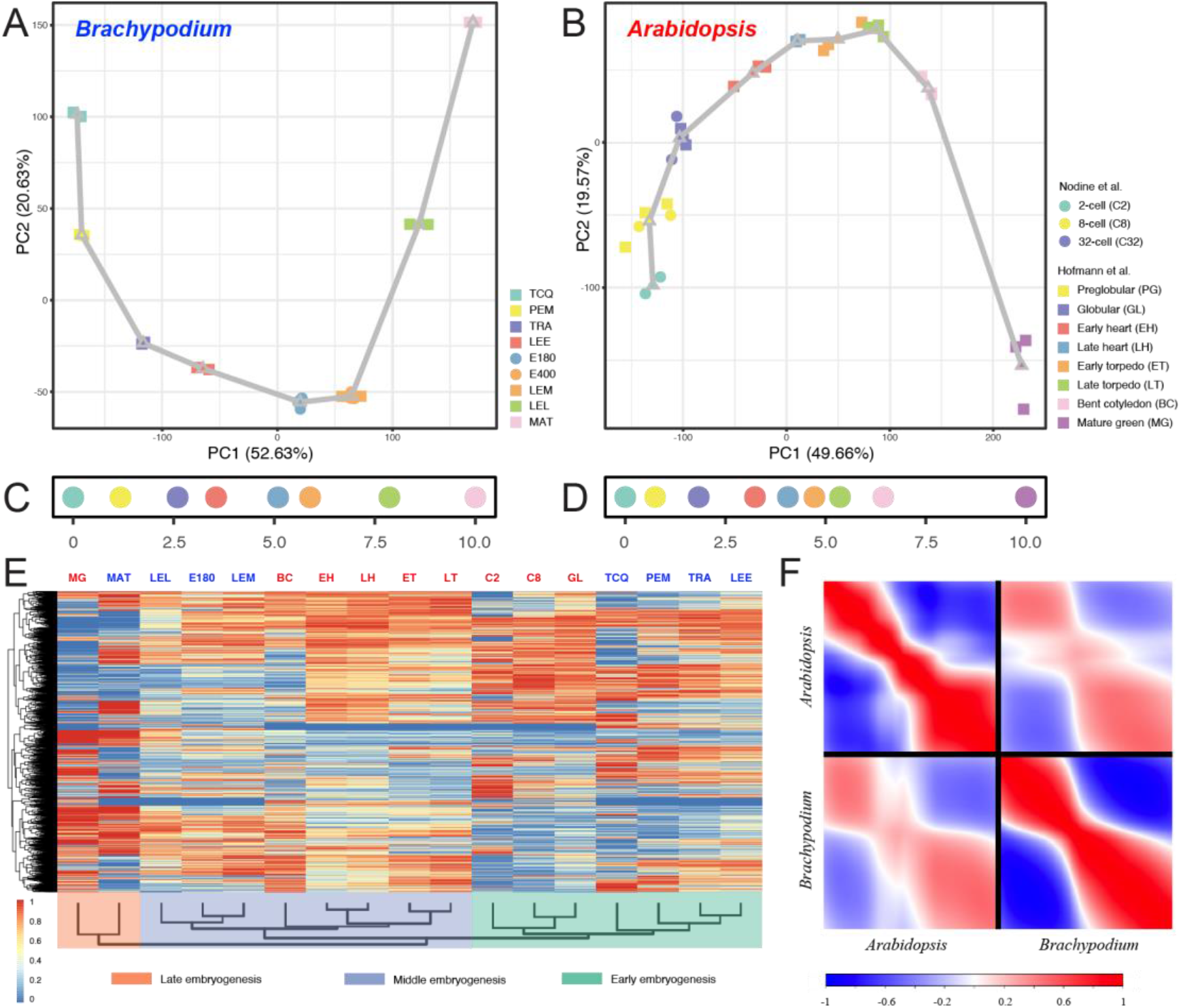
Transcriptional dynamics of *B. distachyon* and *A. thaliana* embryogenesis. All of the differentially expressed genes were used in the PCA plots to stratify embryonic transcriptomes across stages in *Brachypodium* (a) and *Arabidopsis* (b). Each developmental stage was represented using the centroid location in each PCA plot and adjacent centroids were linked using straight lines, forming an expression trajectory. Then, the rank and distance along the developmental trajectory was used to calculate a Developmental Time Units (DTU) value, scaled from 0.0 to 10.0, as a pseudotime metric in each species (c, d). (e) The hierarchical clustering of the relative expression data of orthologous genes between *Brachypodium* and *Arabidopsis* shows broad clustering according to embryogenesis phase (early/middle/late). Stage abbreviations are the same as in the key to (a) and (b). (f) The intra- and interspecific comparisons of embryonic developmental transcriptomes between *Brachypodium* and *Arabidopsis*. The successive stages of embryogenesis (DTU plots from panels c and d) are plotted on both axes of each plot. Species are indicated on each axis. Pearson correlation: positive, red; no correlation, white; negative, blue.

To relate developmental progression between *Arabidopsis* and *Brachypodium* embryogenesis, we calculated a pseudotime metric (Leiboff and Hake 2019), namely developmental time units (DTUs), to reconstruct the molecular ontogenies for each species using the expression trajectory information in their respective PCA plots (Fig. 3C, D). Then, to determine whether individual *Brachypodium* embryo developmental stages can be matched with comparable *Arabidopsis* stages, we performed a hierarchical clustering based on the relative expression data of orthologous genes. This analysis showed that transcriptomes of classes of embryo stages are more similar between species than with different stage classes within species. While the embryo stages between these two species cannot be directly matched, they can be roughly classified into three distinct developmental phases, i.e., early, middle, and late embryogenesis, irrespective of plant species (Fig. 3E). Within each developmental phase, tissues are clustered by species rather than stages, suggesting obvious species-specific transcriptome signatures. Furthermore, intraspecific embryo transcriptome comparisons also reveal that early and late embryogenesis are more conserved between species, and these are separated by a phase of dramatic differences in gene expression (Fig. 3F).

### Conserved and diverged functions during embryo stages across angiosperms

To study the deep phase conservation and divergence in terms of gene expression between *Arabidopsis* and *Brachypodium* embryo development, we mapped thee expression profile of each gene to one of the developmental phases, i.e., early, middle, and late embryogenesis, to identify phase-specific genes of which expression was restricted to one of these three developmental phases (Fig. 4A-D, I-L). We next compared the patterns of orthologous genes between the two species, and found that phase-specificity of orthologues was much more prominent in early and late embryogenesis than in middle embryogenesis (Fig. 4E-H), consistent with the results of the interspecific transcriptome comparison (Fig. 3F). Gene Ontology (GO) enrichment analysis indicates that the early phase is enriched for ribosome, translation, and DNA replication which are associated with cell growth and proliferation (Fig. 4M, N & Supplementary Fig. 6). In contrast, the late phase is enriched for various enzyme activities, transporters, and signaling which reflect a cell-type specific status, suggesting that this phase is characterized by genes expressed in differentiated and specialized cells (Fig. 4M, N & Supplementary Fig. 6). Compared to the early and late phases, the transcriptomes of middle embryogenesis of *Arabidopsis* and *Brachypodium* are less correlated, with only few overlapped phase-specific orthologous genes (Fig. 3F and Fig. 4G). Those limited gene sets are enriched in chloroplast-related GO functional terms (Supplementary Fig. 7). Thus, while the mid-embryo development phase is morphologically and transcriptionally divergent between *Arabidopsis* and *Brachypodium*, a common characteristic is active chloroplast development.

**Figure 4.**
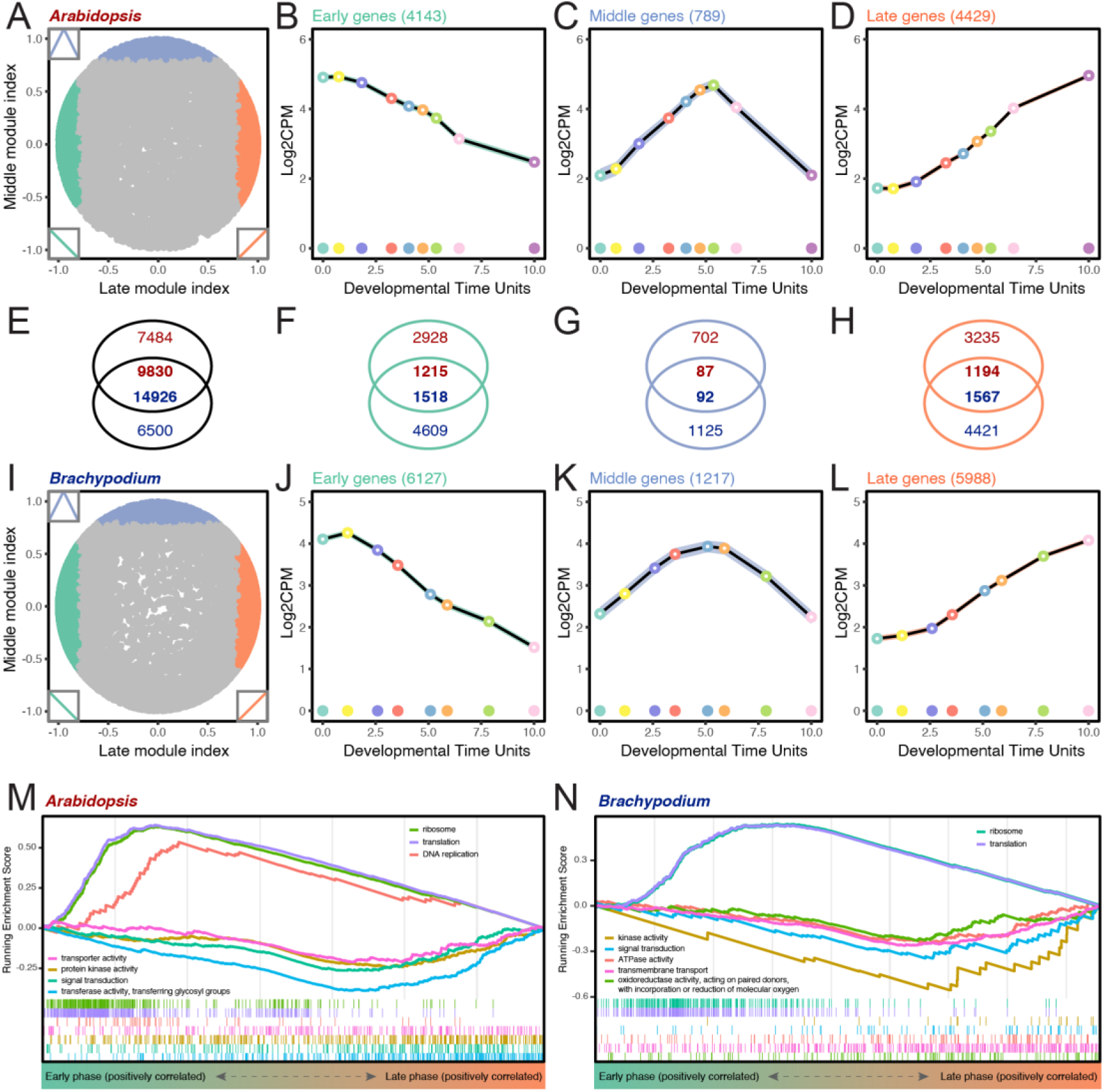
Phase-specific gene expression and functional enrichment. (a, i) Landscapes show the correlations between gene expression profiles and two hypothetical perfect modules: middle (y axis) and late (x axis) modules, during embryogenesis in *Arabidopsis* (a) and *Brachypodium* (i). Each spot corresponds to an *Arabidopsis* (a) or *Brachypodium* (i) gene. The inset graphics show the perfect modules which are used to calculate the correlations. Three sets of phase-specific gene expression are plotted and coloured according to the assigned phases, i.e., early (cyan), middle (blue) and late embryogenesis (orange) in *Arabidopsis* (b-d) and *Brachypodium* (j-l). The number of overlapped orthologous genes for each set is also indicated (e-h). Gene set enrichment analysis (GSEA) shows the gene sets which are highly correlated with early or late phases in *Arabidopsis* (m) and *Brachypodium* (n).

Next, we asked if the middle phase between the early and late embryogenesis is enriched in specific gene families in a species-dependent manner. To achieve this, we first downloaded a collection of gene families of *Arabidopsis* from TAIR (www.arabidopsis.org) and identified homologs in *Brachypodium* by a best blast hit approach. Gene set enrichment analysis (GSEA) indicates a similar result with previous GO enrichment for early and late phases (Supplementary Fig. 8). As for the middle phase, we indeed found species-dependent enriched gene families except for two common families, zinc finger homeodomain (ZF-HD) transcription factor (TF) family and Golden2-like (G2-like) TF family (Supplementary Fig. 9). Interestingly, mid-embryogenesis in *A. thaliana* is enriched for two gene families, *ASYMMETRIC LEAVES2* (*AS2*) and *LATERAL ORGAN BOUNDARIES* (*LOB*), that function in the organ asymmetry and boundary formation (Iwakawa et al. 2002; Semiarti et al. 2001; Shuai et al. 2002), respectively (Supplementary Fig. 9C). Mid-embryogenesis in *B. distachyon* is enriched for the *GROWTH REGULATING FACTOR* (*GRF*) family, which can interact with CUP-DHAPED COTYLEDON (CUC) to promote cotyledon separation in *Arabidopsis* (Lee et al. 2015) (Supplementary Fig. 9D). These results suggest that, while early and late phases are characterized by shared functional programs, the middle phase of embryogenesis is marked by the expression of genes involved in lineage-specific body plans.

### Developmental heterochrony of *B. distachyon* and *A. thaliana* embryogenesis

Among the enriched gene sets in early and late embryogenesis, we identified several TF families. Interestingly, there was a clear difference between *Arabidopsis* and *Brachypodium.* In *Brachypodium*, the GL1 enhancer binding protein (GeBP), MADS-box, Homeobox, CCAAT-HAP5, Alfin-like, MYB, and NAC gene families were all enriched in early and late stages (Supplementary Fig. 8B). None of these were enriched in the early phase of *Arabidopsis* embryogenesis (Supplementary Fig. 8A). Instead, we noticed that the Homeobox TF family was enriched in the middle-to-late phase of *Arabidopsis* embryo development (Supplementary Figs. 8A & 9C). Correlation of gene expression with embryo development further confirmed the expression divergence of this gene family between *Arabidopsis* and *Brachypodium* embryogenesis (Supplementary Fig. 10A, C). The same pattern was observed for the bHLH TF family, although this family was not significantly enriched in early embryogenesis in *Brachypodium* (Supplementary Fig. 10B, D). This finding suggest heterochronic genome-wide expression patterns between these two species for transcription factor families. Given that several members of these two TF family are key regulators of embryo patterning and tissue specification (Ito et al. 2002; Radoeva et al. 2019; Tsuda and Hake 2016), it is possible that the embryo patterning process is heterochronic between these species.

To further dissect heterochrony of early embryogenesis between *Arabidopsis* and *Brachypodium*, we initially mapped all *Brachypodium* genes encompassing a Homeodomain (PF00046 and PF05920) on a gene expression phasegram, which was constructed by sorting the gene expression peak along the DTUs, and compared with that of *Arabidopsis* (Fig. 5A). We found that most *Brachypodium* Homeobox genes, of which the expression is restricted to early embryogenesis, have corresponding *Arabidopsis* homologs that are highly expressed at the middle-to-late stage of embryogenesis (Fig. 5A). Thus, most HD-containing transcription factors that are known to control Arabidopsis embryo development, for which a clear ortholog can be identified in *Brachypodium*, show earlier expression in *Brachypodium* than in *Arabidopsis*. Similar patterns could be observed in other TF families (Supplementary Fig. 11). Thus, if these TF homologs are functionally conserved between the two species, the homologous developmental process may occur earlier in *Brachypodium*.

**Figure 5.**
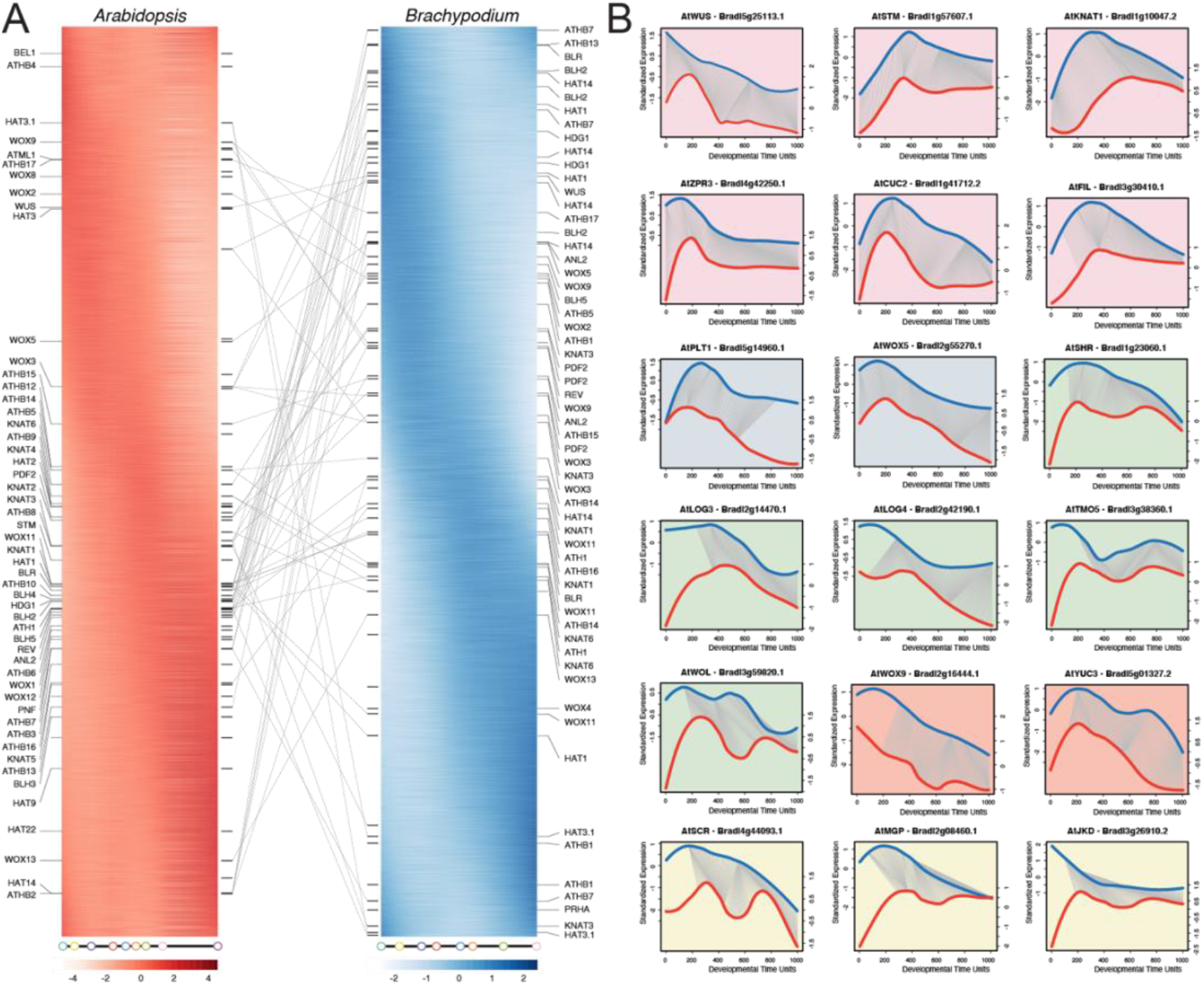
Phased gene expression and dynamic timing warping expression profile alignments. (a) *Arabidopsis* (left, red) and *Brachypodium* (right, blue) homeobox genes are sorted by their expression peaks along embryonic development. All Homeobox family members that are expressed during *Arabidopsis* and *Brachypodium* embryogenesis are mapped to the phasigrams. The lines connect homologous gene pairs of *Arabidopsis* and *Brachypodium*. (b) DTW alignments show comparable expression patterns of cell lineage markers for shoot apical meristem (pink shading; WUS, WUSCHEL; STM, SHOOT MERISTEMLESS; KNAT1, KNOTTED-like 1; ZPR3, LITTLE ZIPPER 3; CUC2, CUP-SHAPED COTYLEDON 2; FIL, FILAMENTOUS FLOWER), quiescent center (blue shading; PLT1, PLETHORA 1; WOX5, WUSCHEL-RELATED HOMEOBOX 5), vascular tissue (green shading; SHR, SHORT ROOT; LOG3, LONELY GUY 3; LOG4, LONELY GUY 4; TMO5, TARGET OF MONOPTEROS 5; WOL, WOODEN LEG), suspensor (orange shading; WOX9, WUSCHEL-RELATED HOMEOBOX 9; YUC3, YUCCA 3), and ground tissue (yellow shading; SCR, SCARECROW; MGP, MAGPIE; JKD, JACKDAW).

### Comparable patterns of activity for developmental regulators across angiosperm embryogenesis

After establishing that there are strong similarities in genome-wide gene expression during early and late embryogenesis, and heterochronic expression of several TF families between *Brachypodium* and *Arabidopsis*, we set out to explore more systematically the correlation between temporal expression patterns of orthologous genes. We performed a dynamic time warping (DTW) expression profile alignment analysis. This analysis will compare the overall expression profiles between two time-series and calculate a DTW distance which is insensitive to local compression and stretches (Giorgino 2009). A low or high DTW distance suggest that a gene pair has a similar or dissimilar expression profile, respectively, between these two time-series datasets (Supplementary Fig. 12A). Comparative enrichment analysis shows that genes with low DTW distances are enriched in genes involved in cell proliferation, such as ‘DNA replication’, ‘cell division’, and ‘THO complex’, of which the expression patterns are correlated with early embryogenesis and the genes annotated with ‘embryo development ending in seed dormancy’ and ‘seed development’ that are normally positively correlated with seed maturation (Supplementary Fig. 12B). These results are consistent with the results above that the fundamental processes in early and late phases during embryogenesis are relatively conserved between *Arabidopsis* and *Brachypodium*.

Although *Arabidopsis* and *Brachypodium* share a conserved developmental program regarding the fundamental processes in early embryogenesis, it is unclear how embryo patterning events, like the establishment of polar axes and the initiation and maintenance of shoot and root apical meristems (SAM and RAM) compare between species. Therefore, we surveyed DTW expression profile alignments and focused mainly on gene pairs of which the *Arabidopsis* homolog is well-known for its expression pattern and role in controlling embryo patterning. Surprisingly, we found that genes involved in SAM specification shared largely comparable temporal expression patterns between *Arabidopsis* and *Brachypodium* (Fig. 5B-F & Supplementary Fig. 13A, B), as well as genes for quiescent center (QC) specification (Fig. 5G, H). For the ground tissue markers, each reached their expression peaks within the early phase, but while most *Arabidopsis* genes retained expression afterwards, their counterparts in *Brachypodium* decreased dramatically afterwards (Fig. 5I-M & Supplementary Fig. 13C, D). Genes that are specifically expressed in the suspensor share a similar temporal expression pattern along with the embryo development between *Arabidopsis* and *Brachypodium* (Fig. 5N, O & Supplementary Fig. 13F). As for vascular tissue specification, gene expression patterns are similar in general, however, *Brachypodium* genes are already highly expressed at the first stage during early embryogenesis compared to their counterparts in *Arabidopsis* (Fig. 5P-S). We thus conclude, on the basis of comparative analysis of patterning genes, that the general progression of patterning is comparable between species. In addition, it appears that many patterning regulators are expressed early in *Brachypodium*, earlier than in *Arabidopsis*, and well before visible signs of organogenesis.

### Auxin activity in early *Brachypodium* embryogenesis

The phytohormone auxin play an important role in early embryogenesis in *Arabidopsis* (Smit and Weijers 2015). In fact, most patterning processes in the *Arabidopsis* embryo appear to depend on auxin response (Moller and Weijers 2009), and interference with synthesis, transport or transcriptional response each cause distinctive patterning defect embryogenesis, and to determine their similarity to *Arabidopsis* orthologs, we determined temporal profiles and DTW analysis of a number of gene families: YUC and TAA/TAR biosynthesis genes, PIN transporters, TIR1/AFB receptors, Aux/IAA repressors and ARF transcription factors (Fig. 6A). For both YUC and TAA/TAR families, we detected expression of at least one member at the earliest stages, although peak of BdTAA/TAR genes expression was later than AtTAA/TAR genes (Fig. 6B, C). Dynamics of PIN gene expression in *Brachypodium* was similar to that in *Arabidopsis*, and suggests early transport activity that persists during embryogenesis (Fig. 6D). TIR1/AFB receptors appear present throughout embryogenesis, and likewise, multiple members of the BdAux/IAA and BdARF families are expressed throughout embryogenesis (Fig. 6E-G). Based on these observations, the predicted capacity to synthesize, transport and respond to auxin in *Brachypodium* is close to that in *Arabidopsis*, and includes the earliest stages. Only few well-characterized auxin-responsive (output) genes have been identified in the *Arabidopsis* embryo (Moller et al. 2017; Schlereth et al. 2010; Vaddepalli et al. 2021). While for some of these, we detected similar temporal profiles of *Brachypodium* orthologs (Fig. 6 & Supplementary Fig. 14), its relevance is unclear.

**Figure 6.**
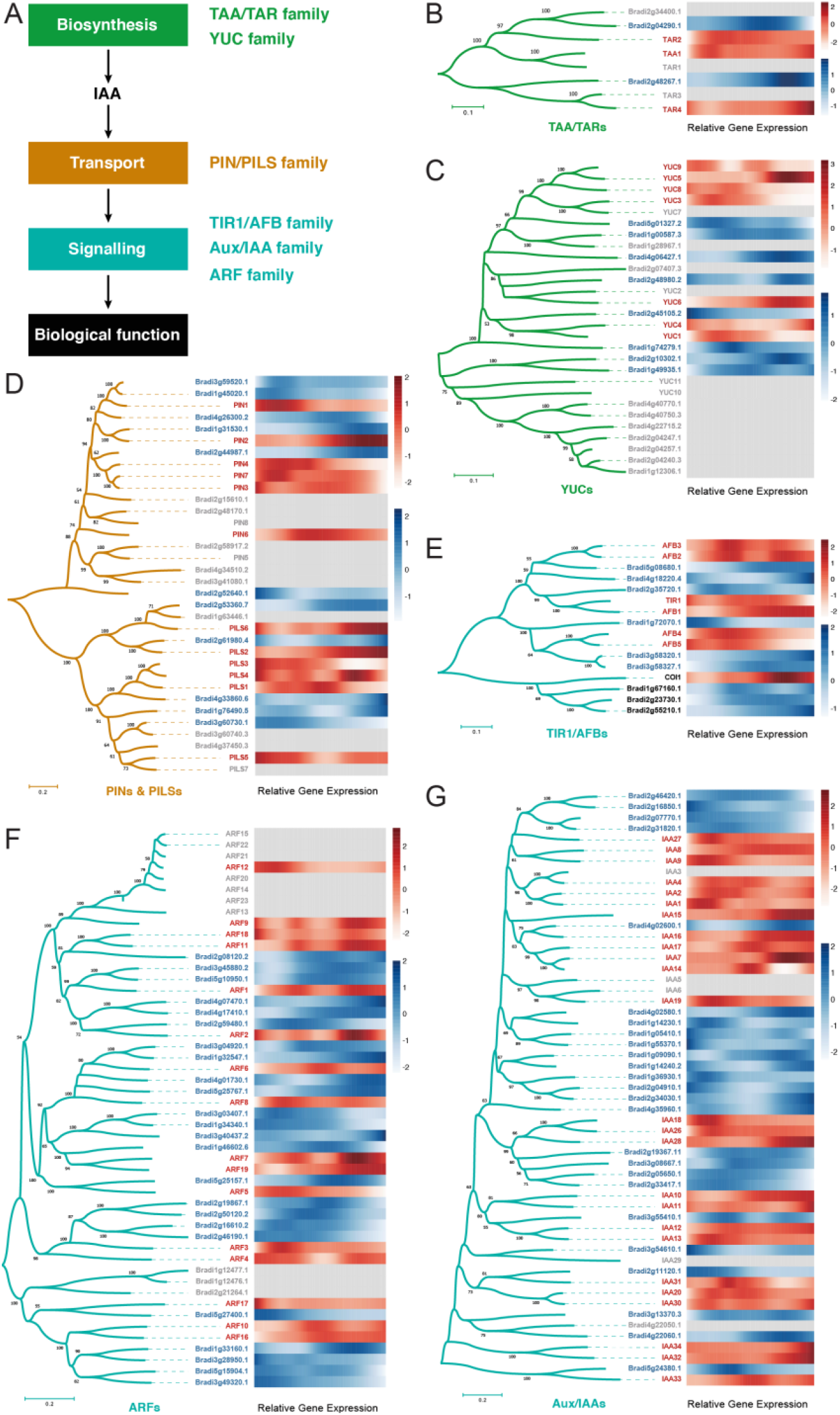
Embryonic expression of auxin-related genes in *Arabidopsis* and *Brachypodium*. (a) A schematic model of gene families that are involved in auxin biosynthesis (TAA/TAR family and YUC family), transport (PIN/PILS family), and signaling (TIR1/AFB family, Aux/IAA family, and ARF family). (b-g) Phylogenetic trees of TAA/TAR (b), YUC (c), PIN1/PILS (d), TIR1/AFB (e), ARF (f) and Aux/IAA (g) gene families were constructed using the NJ method based on genes from *Arabidopsis* (red) and *Brachypodium* (blue). The heatmap of relative gene expression during embryogenesis is shown next to each phylogenetic tree with a white-red (lo-hi)scale for *Arabidopsis* and a white-blue (lo-hi) scale for *Brachypodium*, while genes that were not expressed were shown in grey.

Given the predicted ubiquitous and early auxin activity, we explored the ability of embryos to transport and respond to auxin. To this end, we analyzed the localization of transporters PIN1a, PIN1b and SoPIN1 (*PIN1a-Citrine*, *PINb-Citrine*, and *SoPIN1-Citrine*), as well as expression of the synthetic auxin-responsive promoter DR5 (DR5-RFP) in the developing *Brachypodium* embryo (O’Connor et al. 2014).

As reported by transcriptomics (Fig. 7R), we could indeed detect each PIN protein early during embryogenesis. PIN1a and PIN1b had similar expression patterns (Fig. 7A-M), both were detected in the inner cells of pro-embryo, but PIN1a was more concentrated in the presumed vascular area (Fig. 7A-G) whereas PIN1b was expressed in a broader domain than that of PIN1a (Fig. 7H-M). Furthermore, PIN1a was expressed at a very early stage, which was temporally consistent with the RNA-seq profiling (Fig. 7R), showing a spatially polarized localization towards the one specific cell side (Fig. 7A). Interestingly, soPIN1 had a different expression pattern, which was specifically expressed at the apical domain in the transition stage (Fig. 7N, O) and predominantly expressed at the apical domain afterwards (Fig. 7P, Q). Thus, based on polar localization of PIN proteins, the early *Brachypodium* embryo likely has the capacity to directionally transport auxin.

**Figure 7.**
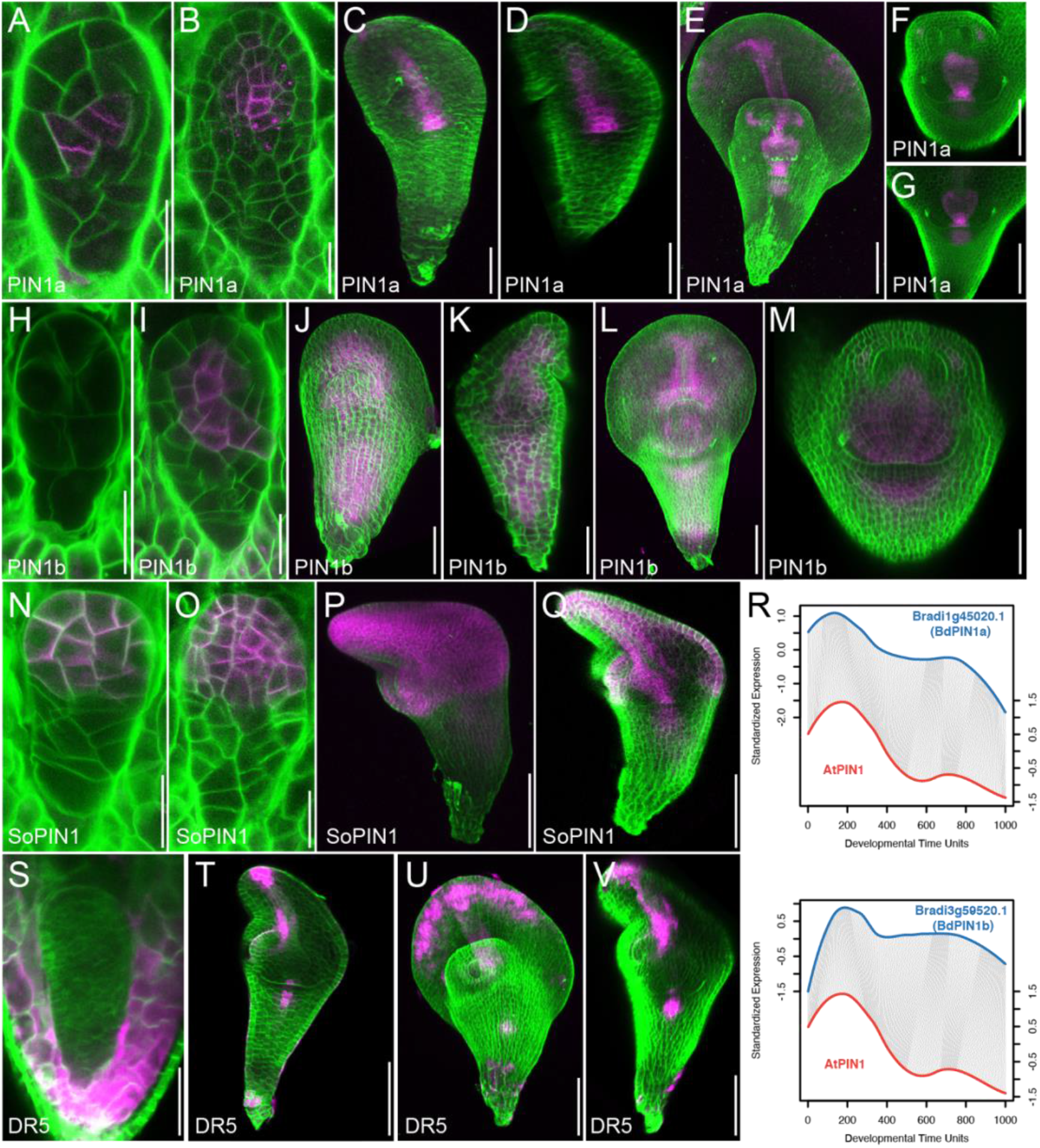
Auxin transport and response during *Brachypodium* embryogenesis. (a-q) Localization of PIN1a-Citrine (a-g), PINb-Citrine (h-m) and SoPIN1-Citrine (n-q) during *Brachypodium* embryogenesis. Magenta color shows PIN protein localization and green color is Renaissance cell wall staining. (r) DTW alignments show comparable temporal expression patterns of PIN1 and SoPIN. (s-v) Expression of DR5-GFP (magenta) in Brachypodium embryos. Green color is Renaissance cell wall staining. Scale bars: 20 μm in (a,b,h,I,n,o), 50 μm in (c,d,j,k,p,q,s), and 100 μm in (e-g,l,m,t-v).

Despite early expression of auxin biosynthesis, transport and response components (Fig. 6) and PIN localization, we did not observe DR5-RFP activity until the transition stage (Fig. 7S-V). At earlier stages, we did detect activity in surrounding maternal tissues (Fig. 7S), similar to patterns observed in *Arabidopsis* and maize (Chen et al. 2014; Robert et al. 2018). From transition stage onward, DR5-RFP was found to be expressed in the root tip, the vasculature, and the tip of the scutellum (Fig. 7T). The auxin signaling at the apical of scutellum was expanded to the entire edge of scutellum in the leaf middle stage (Fig. 7U, V). At these stages, PIN patterns and DR5-RFP activity were in good agreement.

In conclusion, from transition stage onward, the *Brachypodium* embryo is marked by prominent auxin transport and response, which align well with the establishment of vascular tissue and root. At earlier stages, transcriptome analysis predicts extensive auxin activity, but the only available reporter for response could not confirm this activity.

## Discussion

Many fundamental questions related to embryo development in monocots, and by extension, to the conserved and divergent properties between monocots and dicots, have remained unanswered. Here, we have investigated the cellular patterns and gene expression of *Brachypodium* embryogenesis to address such questions. First, we analyzed the pattern of divisions in detail to address the question of whether there is regularity in early divisions. In *Arabidopsis*, most divisions in the embryo are essentially invariant, leading near-complete predictability of the pattern formation process (Mansfield and Briarty 1991). However, this regularity is not shared with many other plants, and descriptions of maize and rice embryogenesis suggest that monocot embryos do not follow a strictly defined pattern of divisions (Chen et al. 2014; Itoh et al. 2005). Through segmentation of cells in early embryos, we find that the first two rounds of cell division in the *Brachypodium* embryo show regularity in both being asymmetric. These two division rounds set apart two small cells from which the central embryo axis likely develops. Later divisions are not strictly controlled, but do further elaborate the early pattern. This finding suggests that there may in fact be a very early pattern formation step in which a domain with distinct developmental fate is defined. Where intuitively, based on morphology, one would perhaps expect pattern formation to be delayed in *Brachypodium*. Compared to *Arabidopsis*, it may in fact commence early.

Through developing a transcriptome resource of the successive embryo stages, we were able to address this question more directly. Given the long time that has passed since the lineages giving rise to *Arabidopsis* and *Brachypodium* split from their last common ancestor, there is no simple orthology relationship between *Arabidopsis* and *Brachypodium* genes, which makes it difficult to infer developmental progression using the expression pattern of known *Arabidopsis* regulators. We therefore first asked if the global progression of embryogenesis is conserved between species. We found that it is, and that particularly the early and late phases of embryogenesis share common transcriptomic features. This suggests, perhaps unsurprisingly, that the physiological and functional processes that mark early and late embryogenesis are conserved across angiosperms. At the same time, the middle phase of embryogenesis showed little global similarity between species, which correlates with the vastly different morphologies observed. Nonetheless, when exploring the temporal expression profiles of the most similar among co-orthologs, and gene family members of known *Arabidopsis* developmental makers and regulators, we did find that their patterns were in fact similar. This is in itself interesting, because it suggests that the timing of expression of developmental genes can be uncoupled from their evidently different spatial patterns, that dictate species-specific morphologies. At the same time, this finding suggests that regulation of development may share substantial parts of regulatory networks across angiosperms.

A notable observation was that there appears to be a clear heterochrony of embryo development between *Arabidopsis* and *Brachypodium*. Based on the expression patterns of large transcription factor families, and other developmental regulators, patterning may occur earlier in *Brachypodium* than in *Arabidopsis*. This is counterintuitive given the lack of early discernable landmarks of embryo patterning in *Brachypodium*, but does align with the finding that the first divisions in the *Brachypodium* embryo are highly regular. We predict based on these findings that the embryo patterning process occurs early in *Brachypodium*, and expect that future investigation of expression patterns of developmental regulators will shed light on the spatiotemporal establishment of cell fates and their regulation.

In *Arabidopsis*, one such regulator is the plant hormone auxin. This small molecule has been implicated in many aspects om embryo development (Moller and Weijers 2009; Smit and Weijers 2015), but based on analysis of the auxin response reporter DR5 in maize, it is questionable if there is active signaling in early embryonic stages (Chen et al. 2014). As was found in the earliest steps of maize embryogenesis (Chen et al. 2014), we also find expression of all components in auxin biology throughout embryogenesis, but the DR5 reporter was likewise not active until transition stage. Active signaling can only be inferred from the expression of genes that are activated by auxin response, of which very few are known, even in *Arabidopsis*. Thus, also in *Brachypodium*, it remains an open question whether early embryonic stages feature auxin response, and if so, whether it contributes to patterning. We do find that the PIN1 proteins, i.e., BdPIN1a, BdPIN1b, and BdSoPIN1 (O’Connor et al. 2014), are expressed, and polarized early during *Brachypodium* embryogenesis. This suggests that at the very least, all hallmarks for active auxin homeostasis, transport, and response are there, which makes is rather unlikely that this system is not used during early embryogenesis. One notable difference between *Arabidopsis* and *Brachypodium* though, is the absence of the PIN3/4/7 clade in the latter. These PIN proteins are prominently active in the early *Arabidopsis* embryo (Friml et al. 2003), and may endow the dicot embryo with unique regulatory abilities. The reporter used to measure auxin response, DR5, is a direct repeat of a medium-affinity binding site for ARF proteins (Boer et al. 2014; Ulmasov et al. 1997), and it is possible, likely even, that this element topology only reports part of the auxin response system. The use of new, high-affinity binding sites with different topologies (Liao et al. 2015) may help to address the question of whether auxin response contributes to early embryo development.

Lastly, our study provides an expression resource for probing genes activity during *Brachypodium* embryogenesis. After maize (Yi et al. 2019) and wheat (Xiang et al. 2019), it is one of the few such resources in monocots, particularly in a non-domesticated species. We expect that deeper analysis will help provide insights into the unique biology of the monocot embryo.

## Materials and Methods

### Plant materials

*B. distachyon* Bd21 plants were grown in growth chambers under long-day conditions of 16 h of light, 22°C and 8 h of dark, 20°C, with light intensity of 100 to 120 μmol^−2^ s^−1^ (Philips high-output F54T5/835-841 bulbs) for the whole life cycle. Spikelets were emasculated and pollinated at the flowering stage to ensure sufficient and developmentally coordinated grain production for embryo isolation. Embryo isolation was performed as described previously (Xiang et al. 2019; Xiang et al. 2011a). For each embryo sample in early stages of development, ~50 embryos were pooled in each biological replicate sample. For each sample in late embryo stages, a minimum of 20 embryos were pooled in each biological replicate sample. A minimum of 20 grains were used for pericarp and endosperm isolation in each biological replicate sample.

### Microscopy

Wheat embryos were cleared in chloral hydrate solution (8:1:2, chloral hydrate:glycerol:water, w/v/v) and viewed with a Leica DMR compound microscope with Nomarski optics. Images were captured using a MagnaFire camera (Optronics) and were edited in Adobe Photoshop CS (Xiang et al. 2011b). Scanning electron microscopy was performed as described previously (Venglat et al. 2011; Xiang et al. 2019) for isolated embryos. For the *Brachypodium* grain, longitudinal hand sections through the grain were made prior to submerging the samples in 25 mM PIPES, pH 7.0, containing 2% (v/v) glutaraldehyde for 2 h. After several washes, the samples were fixed in 2% OsO_4_ in 25 mM PIPES for 2 h, washed, and dehydrated in ethanol (30, 50, 70, 95, and three 100% exchanges).

After sample dehydration, substitution to amyl acetate was performed with increasing ratios of amyl acetate to ethanol (spanning 1:3 parts [v/v], 1:1 [v/v], 3:1 [v/v], then two pure amyl acetate exchanges). All solvent exchanges were separated by 15 min. Samples were critical-point dried with solvent-substituted liquid CO_2_ (Polaron E3000 Series II), mounted on aluminum specimen stubs with conductive carbon glue (Ted Pella), and rotary coated with 10 nm of gold (Edwards S150B sputter coater). Imaging was performed with a 3-kV accelerating voltage, 10-μA current, and 12.2-mm working distance on a Field Emission scanning electron microscope (Hitachi SU8010).

### RNA isolation and antisense RNA amplification

Total RNA was extracted from embryo, endosperm, and pericarp of different developmental stages following the protocol provided by the RNAqueous-Micro kit (Ambion catalog number 1927). The quantity of RNA isolated from early stage embryos was insufficient for library preparation for RNA-seq experiments. Therefore, the mRNA from all stages was amplified and the antisense RNA (aRNA) was used for RNA-seq analysis. The mRNA amplification was conducted according to the protocol provided in the MessageAmp aRNA kit (Ambion catalog number 1750).

### Transcriptome analyses

For RNA-seq profile analysis, we prepared Illumina mRNA-seq libraries using the TruSeq RNA kit (version 1, rev A). Libraries were prepared with aRNA according to the manufacturer’s instructions. For HiSeq 2000 sequencing, four libraries were pooled per sequencing lane. After quality control, read filtering and base correction for the raw read data, we used the clean read data to quantify the expression of representative gene model, the JGI v3.1 annotation of *B. distachyon* Bd21 downloaded from the Phytozome database (http://phytozome.jgi.doe.gov/), using Salmon version 0.13.0 in mapping-based mode with mapping validation (Patro et al. 2017). Read counts were used as the input for differential expression analysis using the Bioconductor package edgeR version 3.24.3 (Robinson et al. 2010). For the time-course data analysis, one-way analysis of variance (ANOVA)-like testing was performed using the glmQLFTest function in edgeR with an FDR cutoff of 0.05.

Sample psuedotime indexing was performed as described in previous studies with some modifications (Leiboff and Hake 2019). Specifically, all of the differentially expressed genes were used to separate samples on a PCA plot for each species. Then, each developmental stage was assigned a location using the centroid value and adjacent centroids were linked using straight lines, producing an expression trajectory. Finally, the rank and distance along the developmental trajectory was used to calculate a Developmental Time Units (DTU) value, scaled from 0.0 to 10.0.

For temporal phased gene expression profiling (Levin et al. 2016), we standardized the logCPM profile by subtracting the mean and dividing by the standard deviation. Then, we calculated the fitted curve for each gene and interpolated the curve into 1,000 points along its psuedotime metric for smooth and continuous comparisons. Next, we calculated the correlation between each gene’s expression profile and two perfect modules, late and middle modules. Since the expression profile was standardized, genes formed a circle as shown in Fig. 4, with x and y axis represented the correlation with the late and middle module, respectively.

To generate a phasegram (Leiboff and Hake 2019; Levin et al. 2016), we perform a PCA analysis for separating genes based on the standardized expression data. The first two components, component 1 and component 2, was used to draw the PCA plot. As the expression dataset were standardized, the genes form a circle. Then, the atan2 function was used to order genes based on the time of expression peak, producing the phasegrams as shown in Fig. 5A.

Dynamic time warping (DTW) was performed on *Arabidopsis*-*Brachypodium* gene pairs, as determined by best blast hit approach, using the standardized expression profiles as described above and the R package dtw v1.22-3 (Giorgino 2009). GO and KEGG enrichment analyses were performed using the Bioconductor package clusterProfiler version 3.10.14 (Yu et al. 2012). Heatmaps were drawn by using the R package pheatmap version 1.0.12.

### Imaging of Auxin reporters

Reporter lines were obtained from O'Connor *et al.* (O’Connor et al. 2014). Tissue was prepared as following: remove the lemma of the spikelet and outer layer of the ovule and excise the exposed young seed from the tip of spikelet (200-400 μm). The excised tissue was then fixed and cleared according to the published method (Ursache et al. 2018). We used a SP5 upright confocal microscope (Leica) to perform all imaging analyses. Excitation wave lengths for different fluorescence markers were as following: UV laser with 405 nm for Renaissance and 514 nm for mCitrine. Images were processed using LasX (Leica) or MorphGraphX software (Barbier de Reuille et al. 2015), and visualized by Photoshop (Adobe).

### 3D segmentation

Tissue was prepared and imaged as above. The Z-stack depth is 0.13 μm. 3D segmentation was performed as described in previous studies (Barbier de Reuille et al. 2015).

## Availability of data and material

Raw reads have been deposited in the NCBI Genome Expression Omnibus under accession number GSE168154.

## Code availability

Not applicable.

## Authors’ contributions

D.W. was the lead investigator of this research program. Z.Z., Z.H., and D.W. designed the experiments and coordinated the project. Z.Z., P.V., D.X., P.G., and R.D. performed the experimental work and collected samples. Z.H. and J.C. performed the comparative transcriptome analyses. Z.H., Z.Z., and D.W. wrote and edited most of the manuscript. All authors have read and approved the final manuscript.

## Acknowledgements

We are grateful to Devin O’Connor for sharing the PIN and DR5 reporter lines, Christian Hardtke for sharing Bd-21 seeds and Sumanth Mutte for helpful discussions.

