## Supplemental Figures 1-14 for "Conserved, divergent and heterochronic gene expression during *Brachypodium* and *Arabidopsis* embryo development"

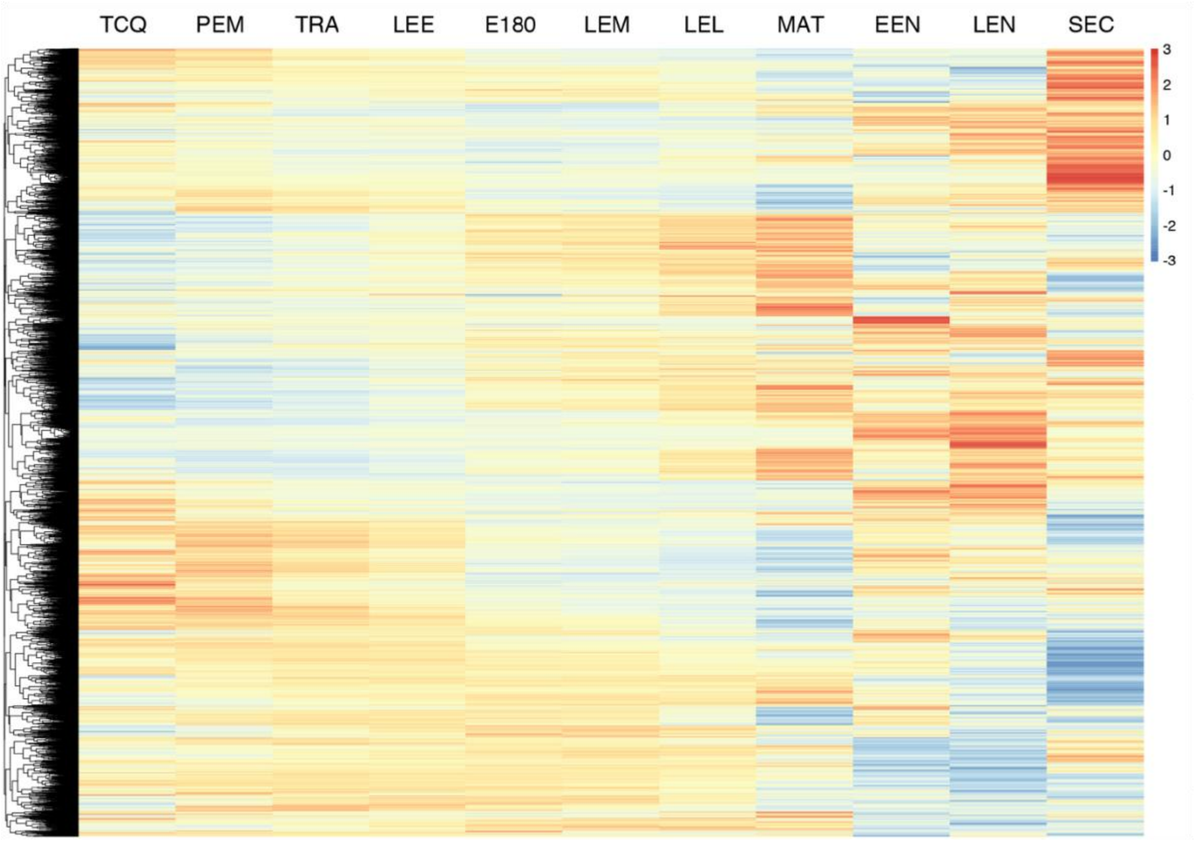


**Supplementary Figure 1. Transcriptome profiling of *B. distachyon* embryogenesis.** Hierarchical clustering of differential gene expression. Embryonic tissues cover eight successive developmental stages (TCQ, PEM, TRA, LEE, E180, LEM, LEL and MAT), two endosperm tissues correspond to embryonic TRA stage (EEN) and LEL stage (LEN), respectively, and the seed coat (SEC) corresponds to the embryonic LEL stage. Color scale show relative expression.

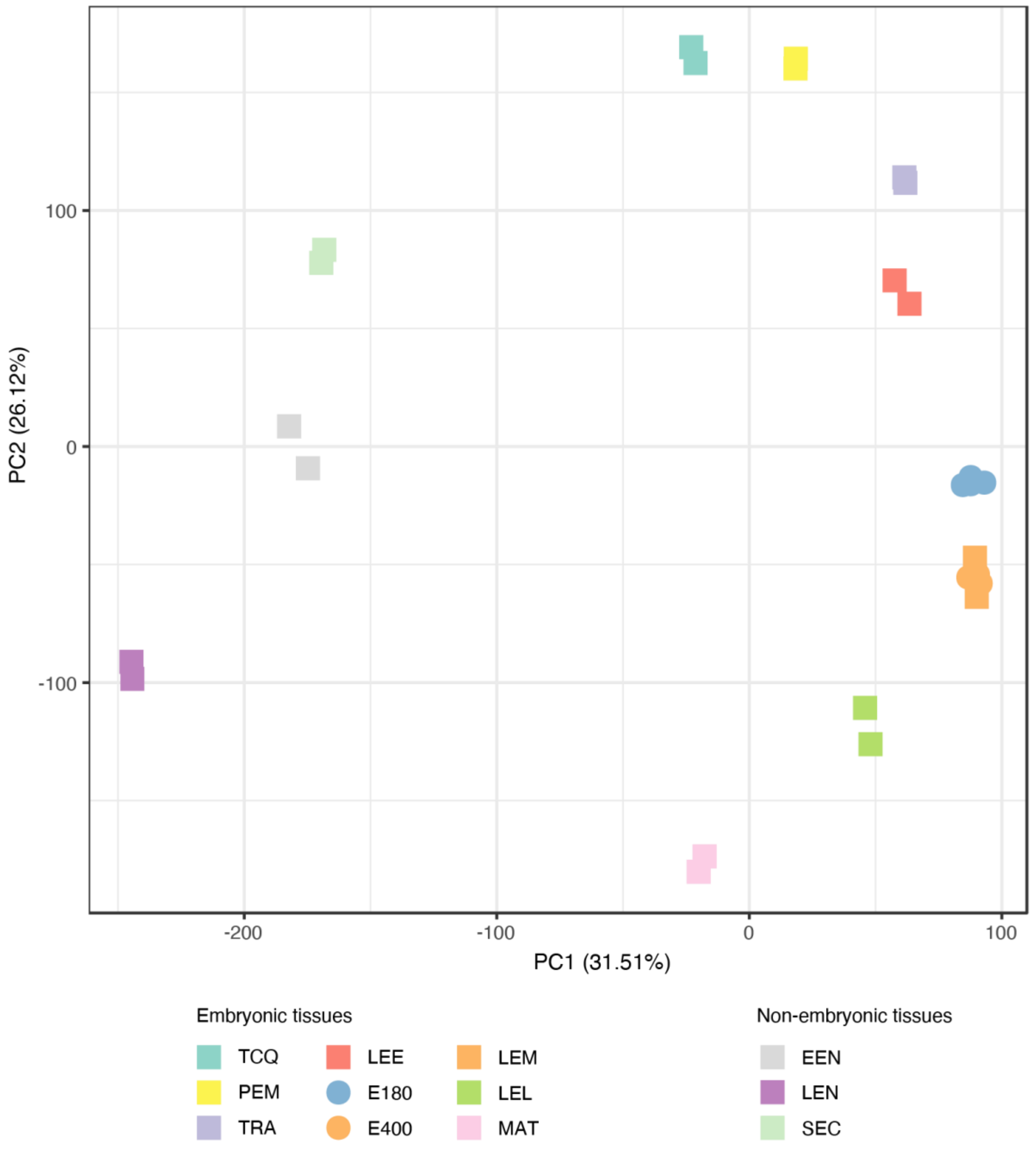


**Supplementary Figure 2. Principal Component Analysis of Brachypodium transcriptome samples.** PCA plot depicting the variance among replicates of each embryo sample. PC1 and PC2 explain 31.51% and 21.62% of the total variance, respectively. Each stage or tissue has two replicates, except for E180 and E400 which both have four replicates. For each sample, replicates are all clustered very closely and separated from all other samples. In addition, there is a clear distinction between embryonic and non-embryonic tissues. Among embryonic tissues, there is clear separation of successive stages through PC1 and PC2.

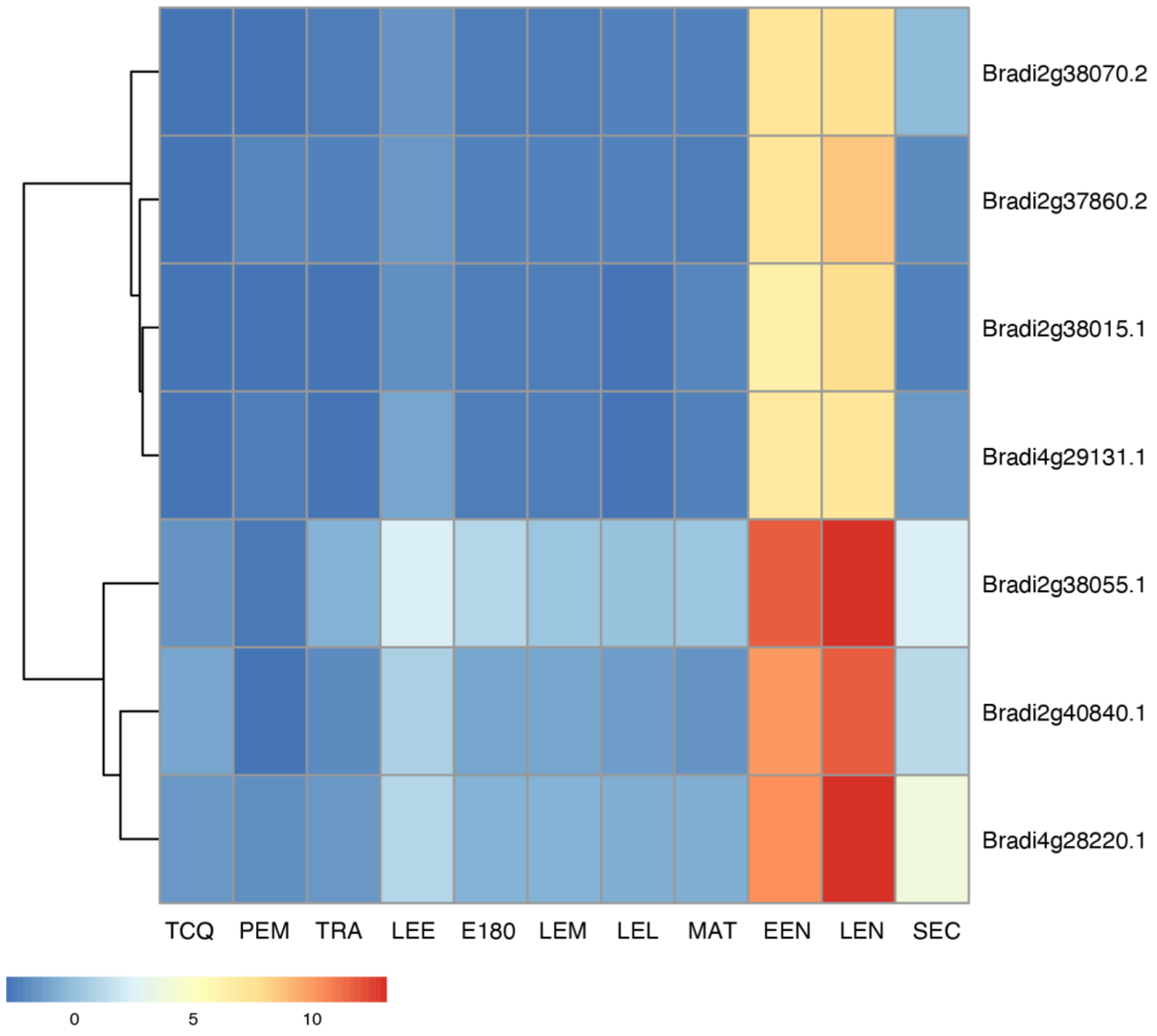


**Supplementary Figure 3. Glutelin gene expression in embryo and non-embryo samples.** The relative expression of 7 Brachypodium glutelin genes is plotted across the 8 embryonic stages, and in the 3 non-embryo samples. Relative expression is shown in color scale.

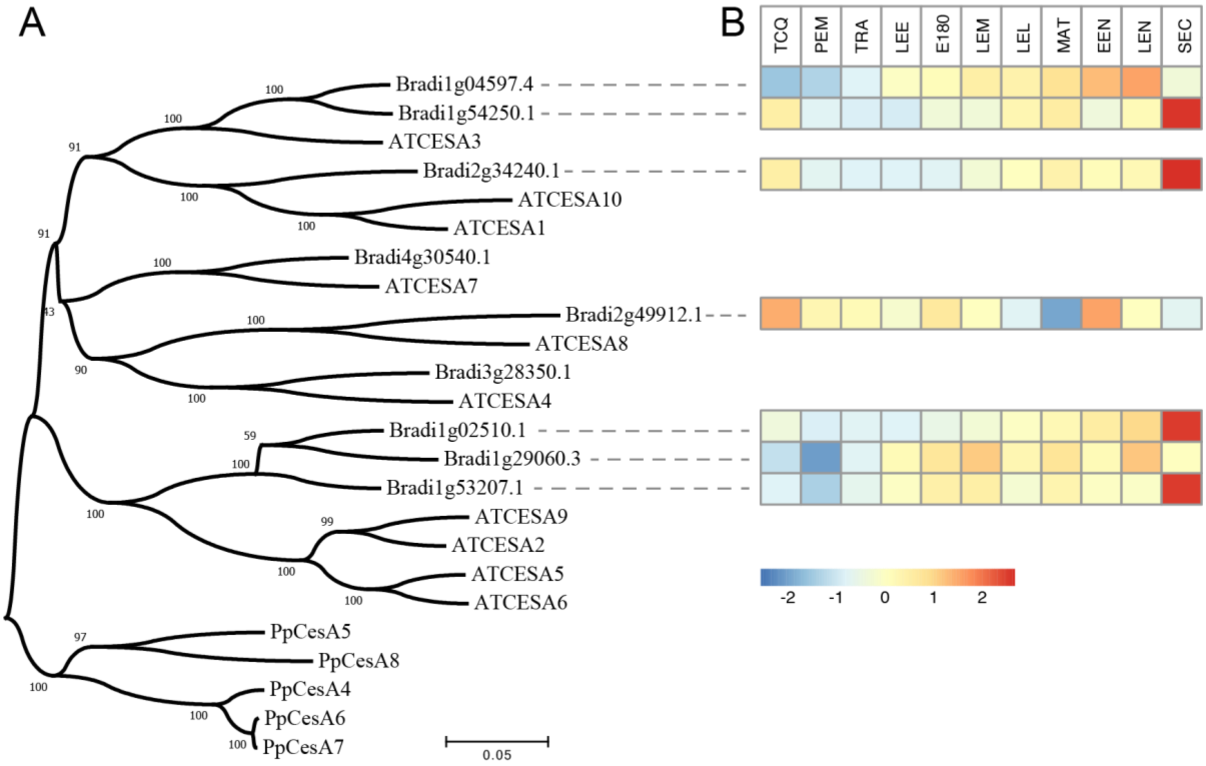


**Supplementary Figure 4.** ***CESA*** **gene expression during** ***Brachypodium*** **embryogenesis.** (a) Phylogenetic tree of *CESA* genes from *Arabidopsis*, *Brachypodium* and *Physcomitrium patens* and (b) heatmap of *CESA* gene expression in *Brachypodium* embryo and non-embryo samples. Relative expression is shown in color scale.

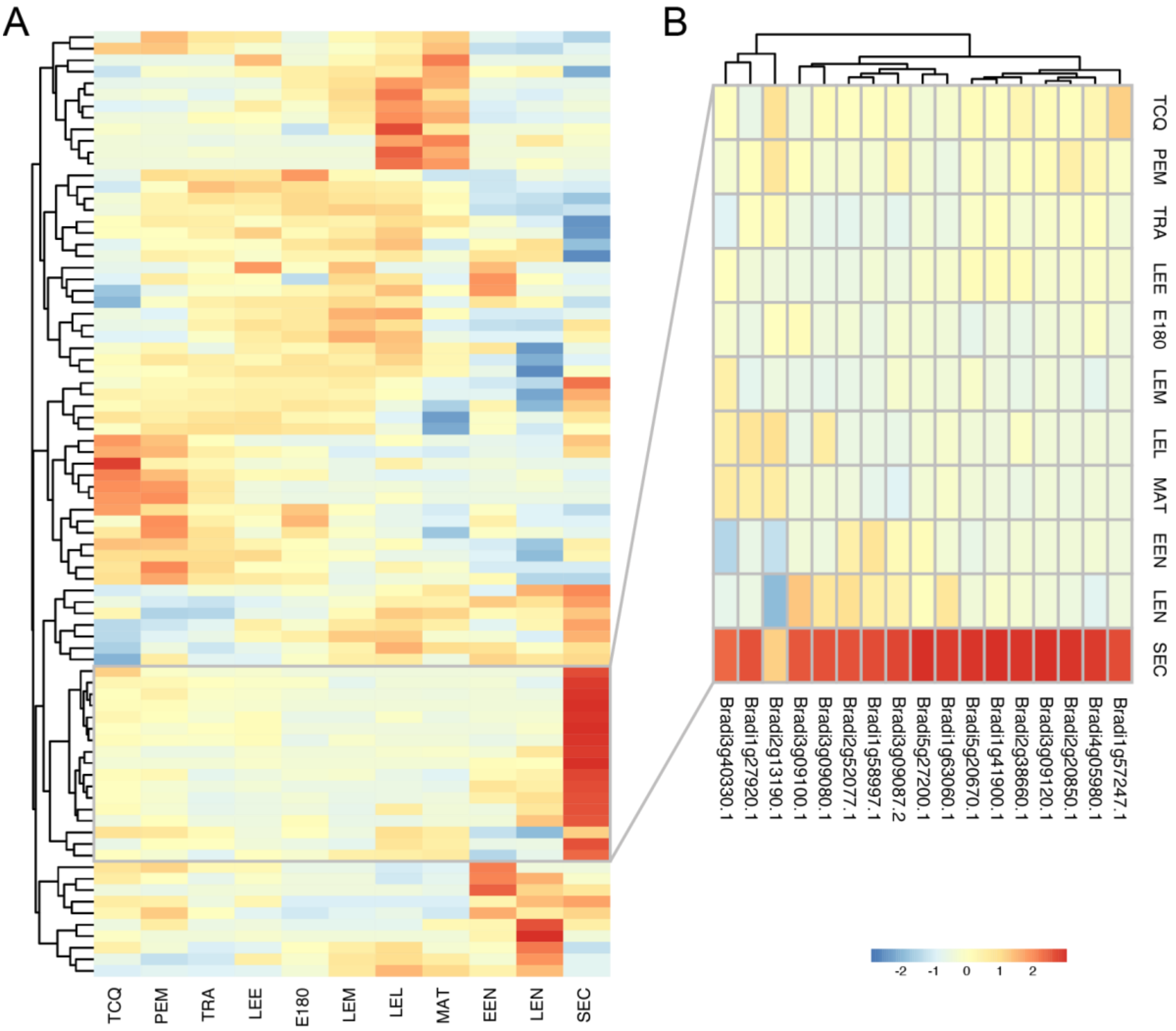


**Supplementary Figure 5. Peroxidase genes expression during *Brachypodium* embryogenesis.** (a) Hierarchical clustering of 82 genes encoding peroxidases that are expressed during *Brachypodium* embryogenesis, along with relative expression in embryo and non-embryo samples. (b) Detailed heatmap of peroxidases that are highly expressed in seed coat samples. Relative expression is shown in color scale.

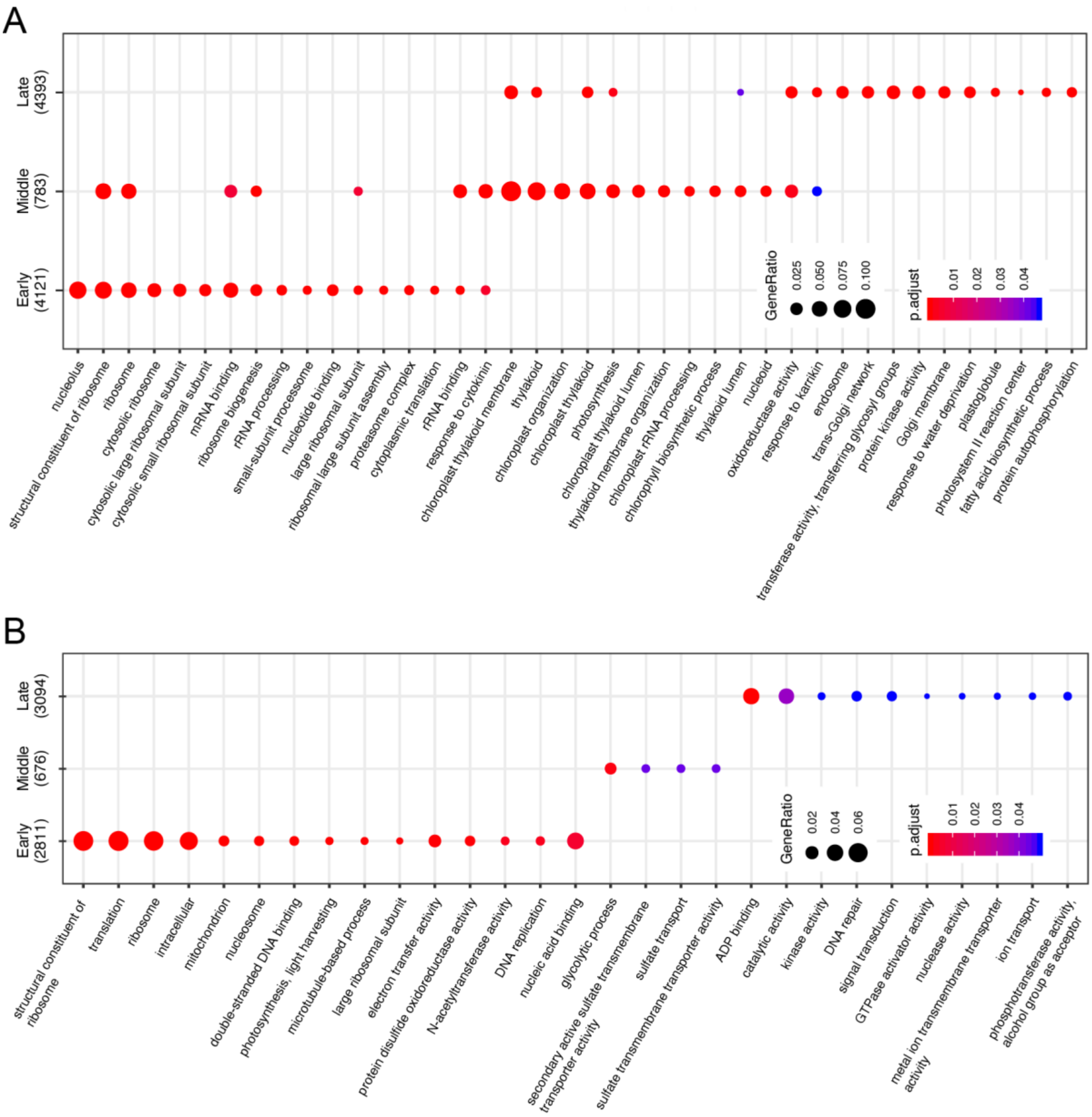


**Supplementary Figure 6. Gene Ontology (GO) enrichment analyses of embryo phases.** Plots show the enrichment (gene ratio indicated by size of circle; adjusted p-value indicated by color scale) of functional annotations in gene sets that are highly expressed during early, middle or late embryogenesis in *Arabidopsis* (a) or *Brachypodium* (b).

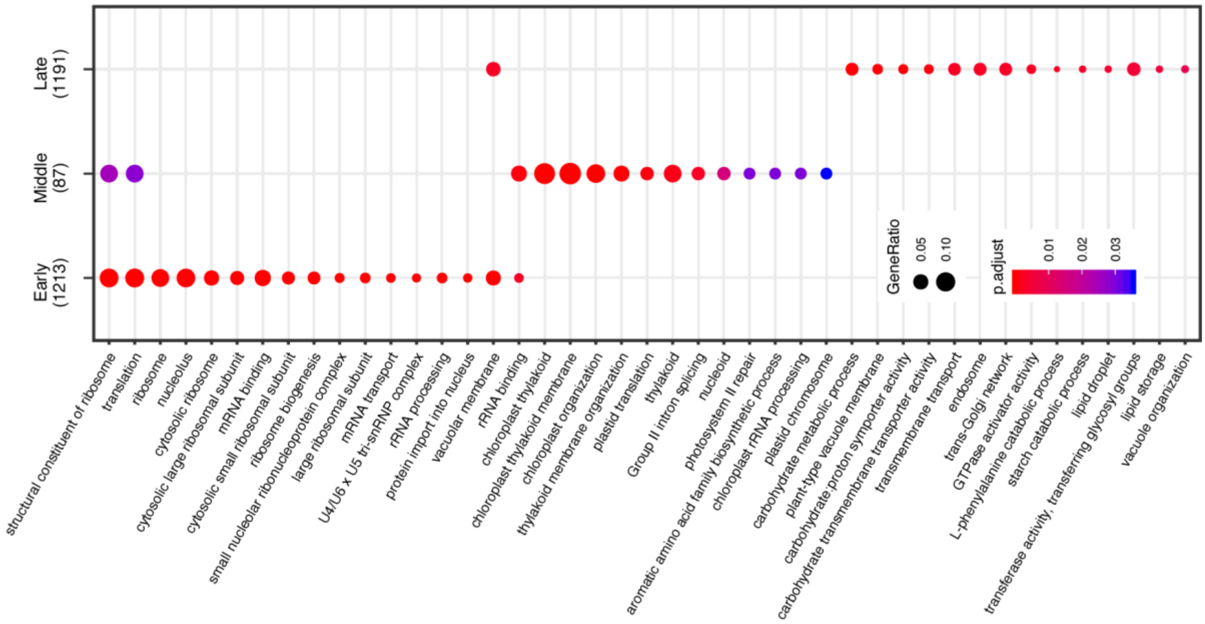


**Supplementary Figure 7. Gene Ontology (GO) enrichment analysis of conserved phase-specific genes.** Plot shows the enrichment (gene ratio indicated by size of circle; adjusted p-value indicated by color scale) of functional annotations in gene sets that are highly expressed during early, middle or late embryogenesis, and for which orthologous gene pairs in *Arabidopsis* and *Brachypodium* share the same phased expression.

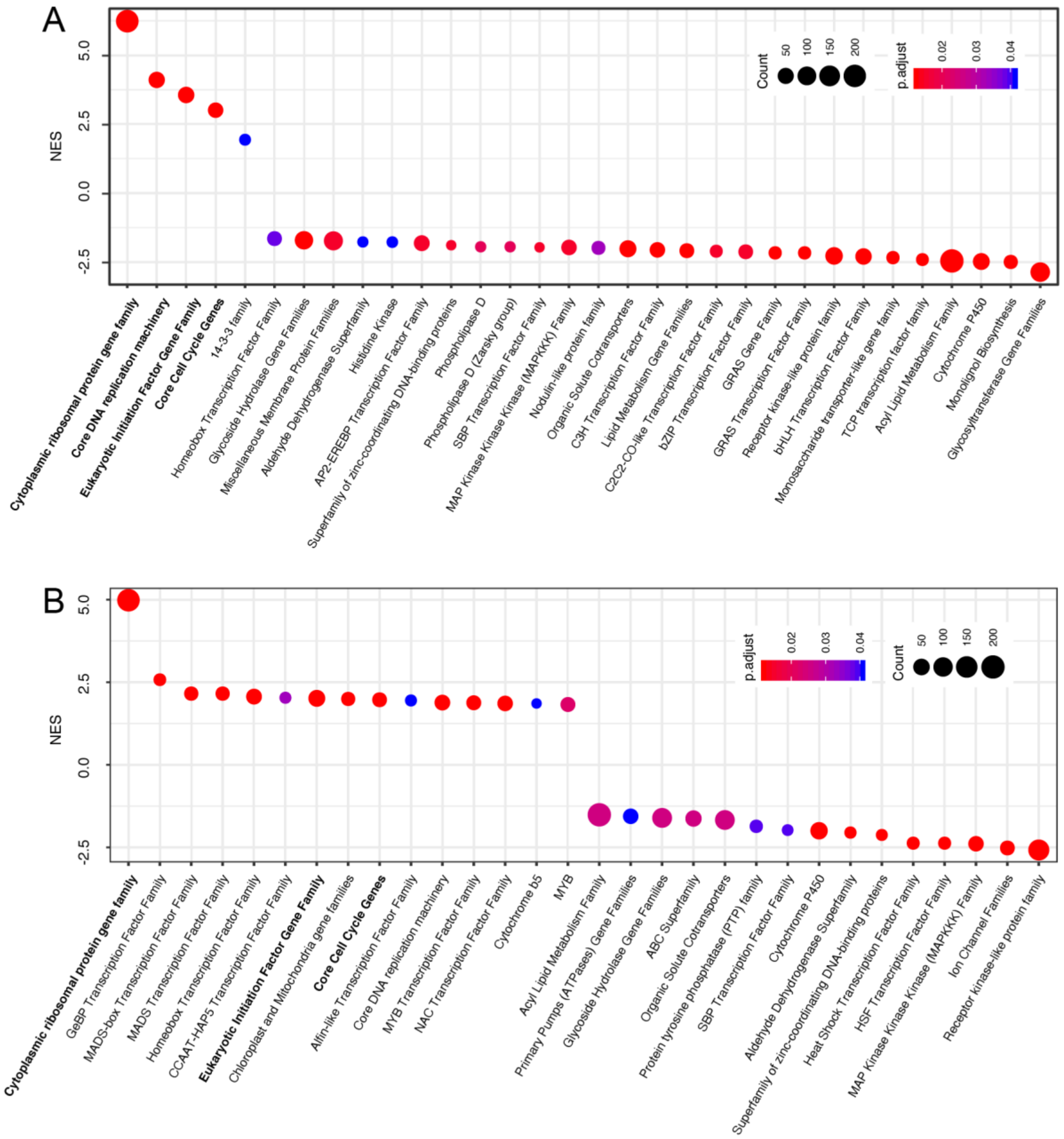


**Supplementary Figure 8. Gene set enrichment analysis (GSEA) of early and late embryonic developmental genes.** Both *Arabidopsis* (a) and *Brachypodium* (b) enriched ribosome, cell cycle, and translation initiation which are related to cell proliferation in the early phase while the late phase is enriched for the genes involved in various enzyme activities, transporters, and transcription factors which are correlated with cell differentiation. The y axis represents for normalized enrichment score (NES).

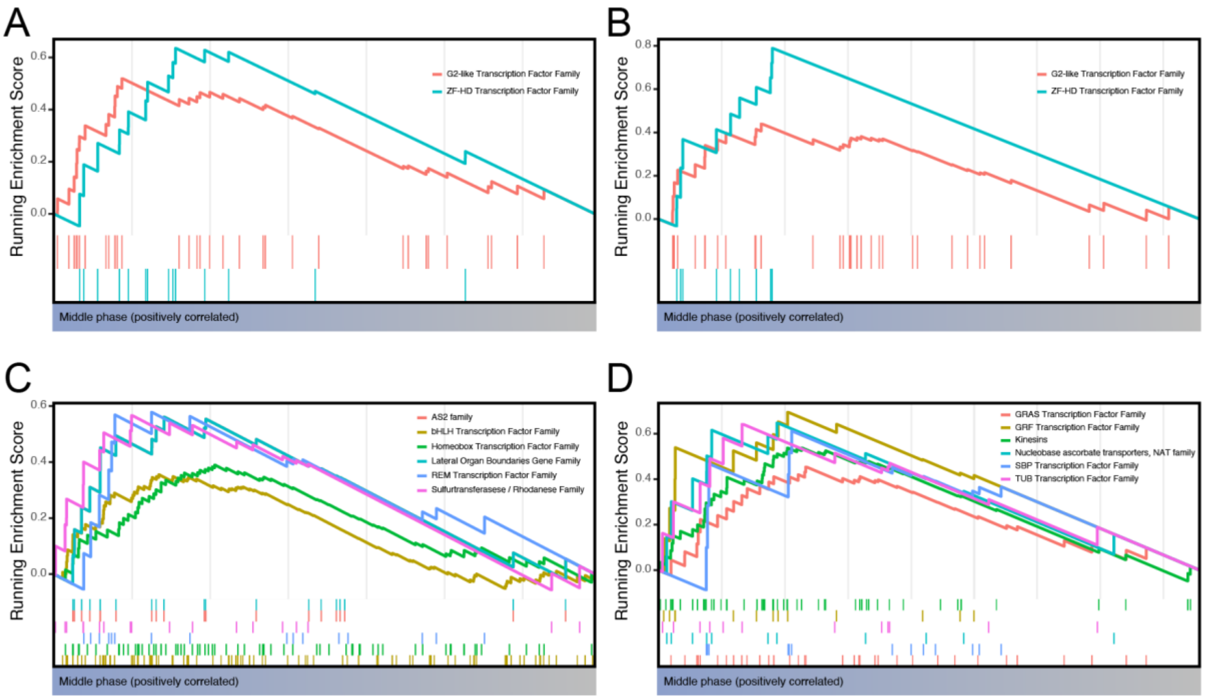


**Supplementary Figure 9. Gene set enrichment analysis (GSEA) of middle embryonic developmental genes.** Both *Arabidopsis* (a) and *Brachypodium* (b) middle phases during embryogenesis are enriched in ZF-HD transcription factor family and G2-like transcription factor family. Except for these two common gene families, the middle phases of *Arabidopsis* (c) and *Brachypodium* (d) are enriched in various but different gene families.

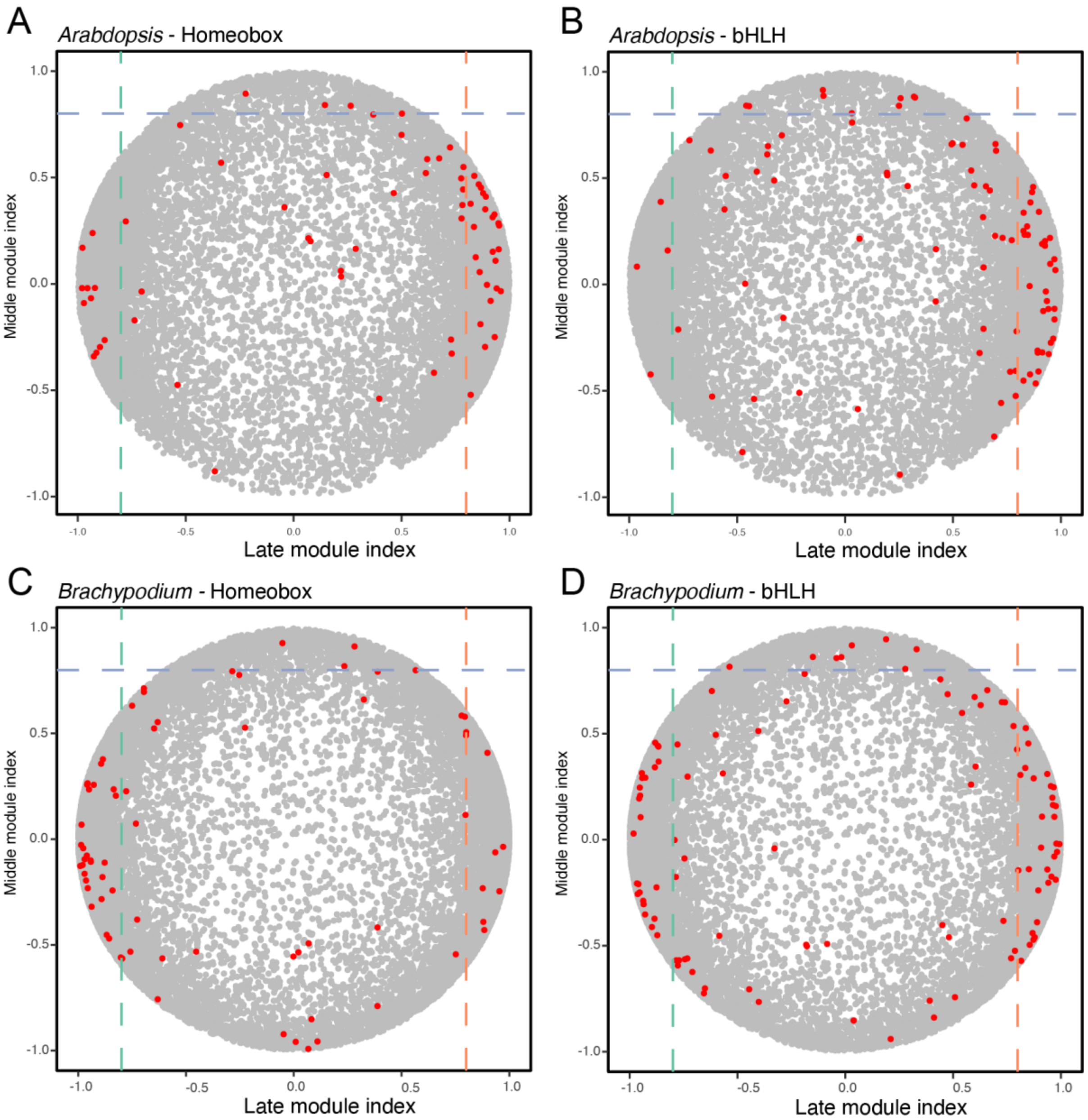


**Supplementary Figure 10. HD and bHLH expression profiles across *Arabidopsis* and *Brachypodium* embryogenesis.** Members of the HD (a,c) or bHLH (b,d) transcription factor families are plotted on the middle/late module index plot for Arabidopsis (a,b) or Brachypodium (c,d). Each TF gene is represented by a red dot.

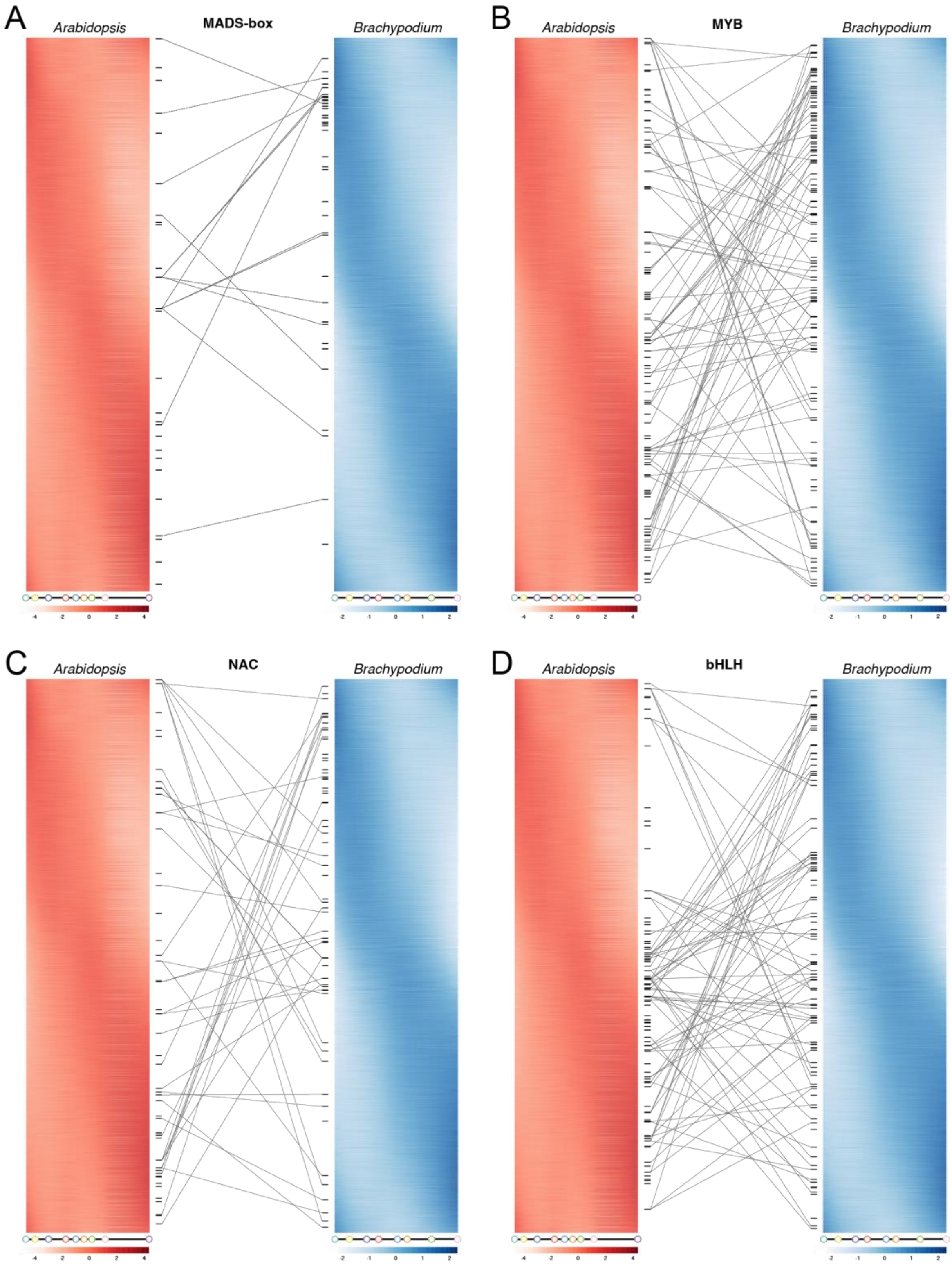


**Supplementary Figure 11. Phased gene expression and gene family mapping.** All members of the *Arabidopsis* (red) and *Brachypodium* (blue) MADS-box (a), MYB (b), NAC (c) and bHLH (d) gene families are sorted by their expression peaks along embryonic development. relative expression is shown in white (low) to red or blue (high) scale, and DTU plots below each phasigram indicate developmental progression. Orthology relationships between Arabidopsis and Brachypodium genes are indicated by connecting lines.

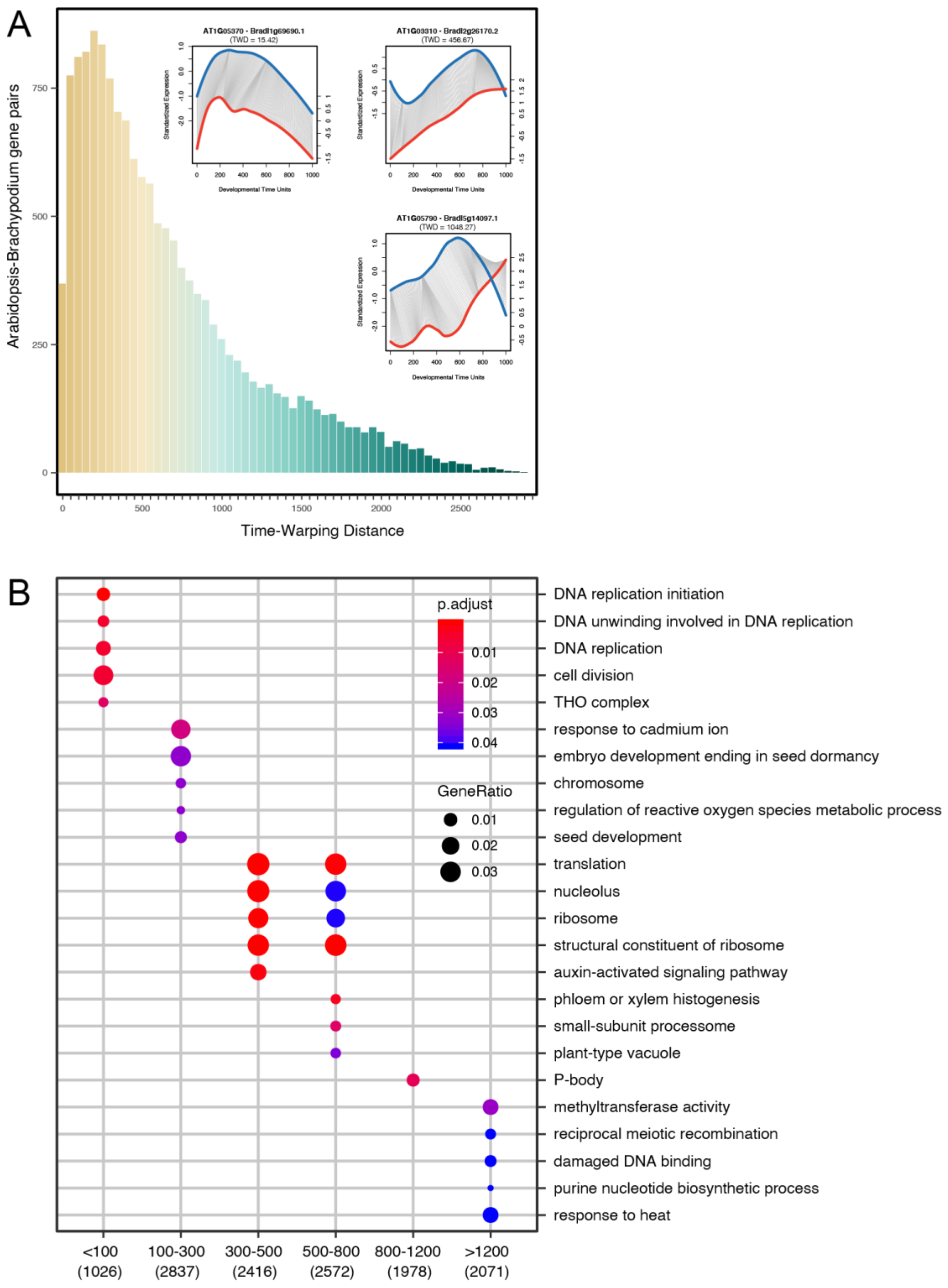


**Supplementary Figure 12. Dynamic time warping analysis of *Arabidopsis* and *Brachypodium* embryogenesis.** (a) Examples of expression time warping plots for orthologous gene pairs between Arabidopsis and Brachypodium throughout embryogenesis (insets; including time warp distance [TWD]), and distribution of TWD distance scores for all *Arabidopsis-Brachypodium* gene pairs. (b) Gene Ontology analysis of gene sets binned by time warping distance score (x-axis). Plot shows the enrichment (gene ratio indicated by size of circle; adjusted p-value indicated by color scale).

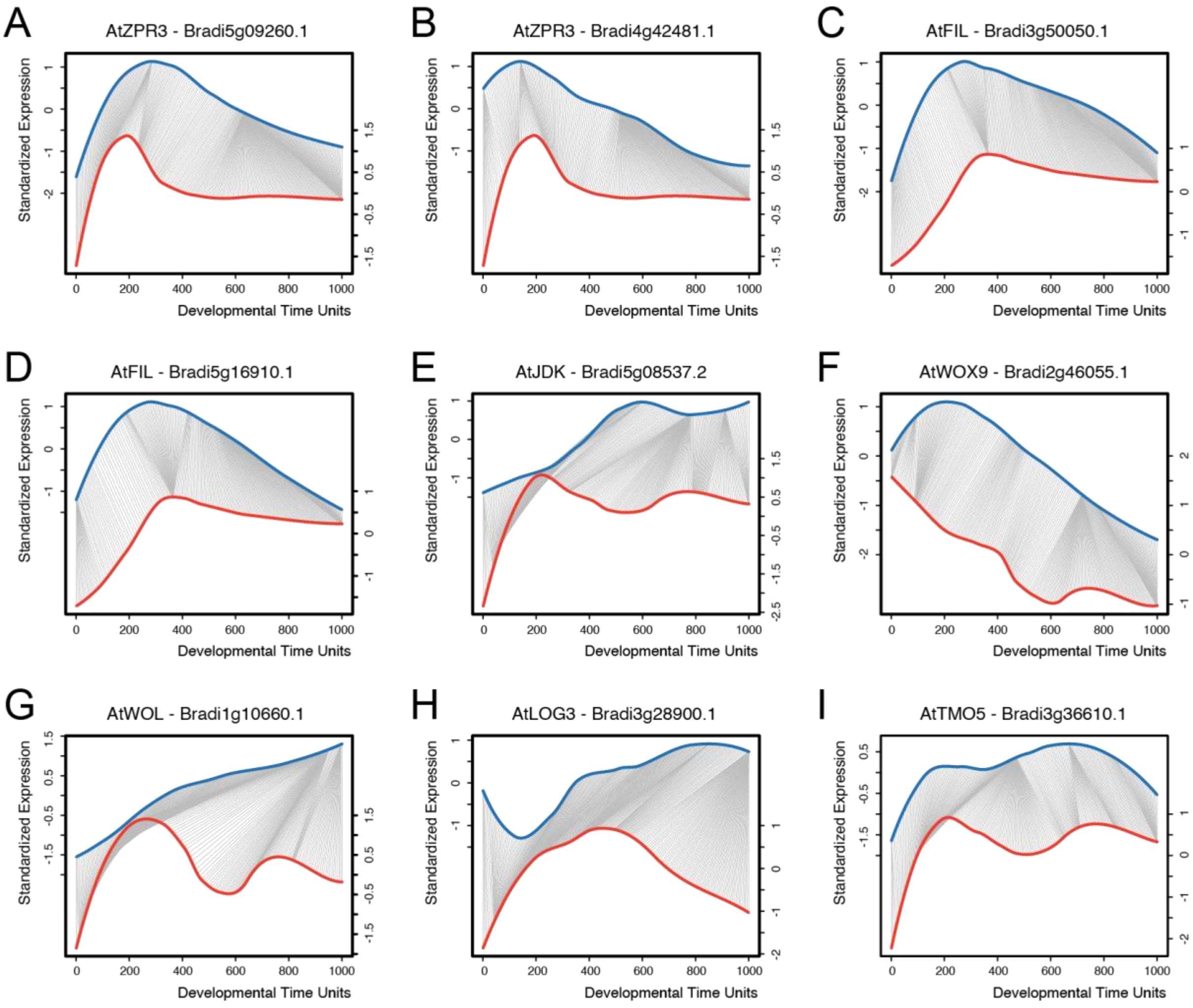


**Supplementary Figure 13. TWD expression profile alignments of gene pairs between *Arabidopsis* and *Brachypodium*.**

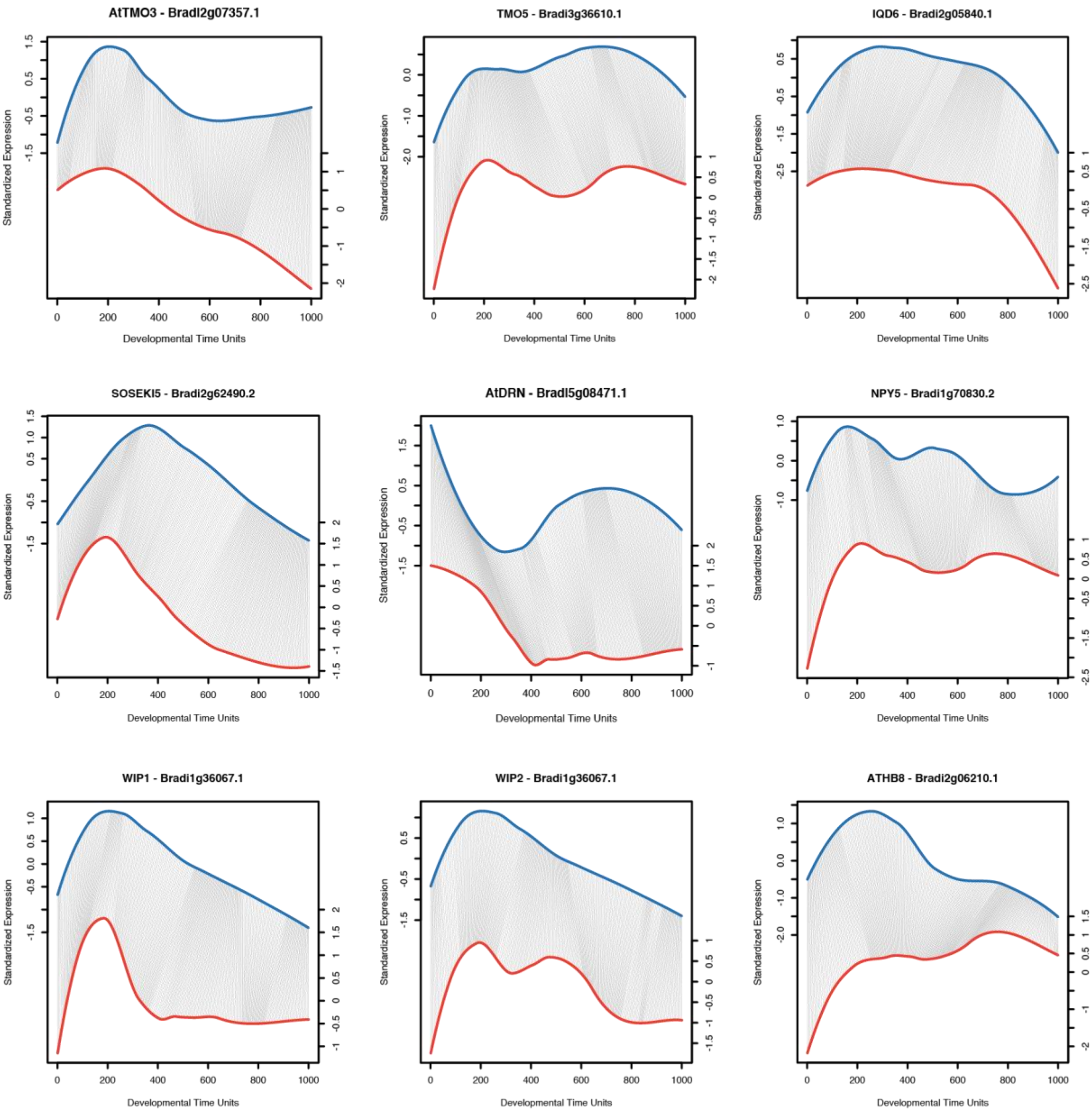


**Supplementary Figure 14. DTW expression profile alignments of *Arabidopsis*-*Brachypodium* gene pairs, of which *Arabidopsis* counterparts are well-characterized auxin-responsive (output) genes during embryogenesis.**
